# LRRK2 G2019S kinase activity triggers neurotoxic NSF aggregation

**DOI:** 10.1101/721266

**Authors:** Francesca Pischedda, Maria Daniela Cirnaru, Luisa Ponzoni, Michele Sandre, Alice Biosa, Maria Perez Carrion, Oriano Marin, Michele Morari, Lifeng Pan, Elisa Greggio, Rina Bandopadhyay, Mariaelvina Sala, Giovanni Piccoli

## Abstract

Parkinson’s disease (PD) is characterized by the progressive degeneration of dopaminergic neurons within the substantia nigra pars compacta and the presence of protein aggregates in surviving neurons. LRRK2 G2019S mutation is one of the major determinants of familial PD cases and leads to late-onset PD with pleomorphic pathology, including alpha-synuclein accumulation and deposition of protein inclusions. We demonstrated that LRRK2 phosphorylates N-ethylmaleimide sensitive factor (NSF). We observed aggregates containing NSF in basal ganglia specimens from G2019S carrier PD patients and in cellular and animal models expressing the LRRK2 G2019S variant. We found that LRRK2 G2019S kinase activity induces the accumulation of NSF in toxic aggregates. Noteworthy, the induction of autophagy cleared NSF aggregation and rescued motor and cognitive impairment observed in aged hG2019S BAC mice. We suggest that LRRK2 G2019S pathological phosphorylation hampers substrate catabolism, thus causing the formation of cytotoxic protein inclusions.

**Highlights:**

- LRRK2 phosphorylates NSF *in vivo*
- NSF aggregates in complementary LRRK2 G2019S models
- LRRK2 G2019S kinase activity induces NSF accumulation in toxic aggregates
- Autophagy induction rescues hG2019S BAC mice motor and cognitive impairment

## Introduction

Parkinson’s disease (PD) is a heterogeneous movement disorder characterized by the progressive degeneration of dopaminergic neurons within the Substantia Nigra pars compacta (SNpc) and the formation of Lewy bodies containing alpha-synuclein in surviving dopaminergic neurons [reviewed in (Obeso et al. 2017)]. Mutations in LRRK2 gene (PARK8; OMIM 609007) are linked to late-onset autosomal dominant PD, accounting for up to 13% of familial PD cases compatible with dominant inheritance (Paisan-Ruiz et al. 2008) and 1 to 2% of sporadic PD patients (J. P. Taylor, Mata, e Farrer 2006). Clinically and pathologically, the features of LRRK2-associated parkinsonism are often indistinguishable from idiopathic PD [reviewed in (Whaley et al. 2006)]. Phylogenetically the LRRK2 kinase domain belongs to the TKL (tyrosine like kinases) and shows high similarity to Mixed lineage kinases (MLKs) (Marin 2006). Like MLKs, LRRK2 is a serine/threonine kinase with no detectable tyrosine kinase activity (West et al. 2007). Disease segregating mutations in LRRK2 have been reported in the kinase domain (G2019S, I2020T), the Roc domain (N1347H, R1441C/G), and in the COR domain (Y1699C) [reviewed in (Berwick et al. 2019)]. However, despite its relevance in PD, the physiological function of LRRK2 and the meaning of PD-linked mutations are yet to be completely understood. The protein inclusions observed in PD patients may be due to the failure of the protein clearance pathways. Intriguingly, experimental evidence suggests that LRRK2 might influence protein clearance acting on both the ubiquitin-proteosome system (Lichtenberg et al. 2011) and autophagy-lysosome pathway [reviewed in (Albanese, Novello, e Morari 2019)]. LRRK2 kinase activity plays a major role in LRRK2 physiological and pathological activity [reviewed in (M. Taylor e Alessi 2020)]. Few LRRK2 substrates have been characterized *in vivo*, including LRRK2 itself (Di Maio et al. 2018) and Rab proteins (Steger et al. 2016). In this study, we described that LRRK2 phosphorylates NSF *in vivo*. Mice expressing hLRRK2 G019S via BAC *(Melrose et al. 2010)* present an age-dependent motor and cognitive impairment together with the deposition of NSF aggregates. At the mechanistic level, LRRK2 phosphorylation triggers the formation of NSF toxic aggregates.

## Results

### hG2019S mice show an age-dependent motor and cognitive impairment

Previous studies described an age-dependent neuropathological and behavioral phenotype in hG2019S BAC (hG2019S) mice (Melrose et al. 2010). We characterized age-matched wild-type and hG2019S male mice at 3, 6, 12, and 18-months in a battery of behavioral tests, including spontaneous motor activity (figure 1A-B), balance beam, pole test, rotarod (figure 1C-G) and novel object recognition (figure 1H). Three-months old wild-type and hG2019S cohorts performed equally in all experiments, except in the rotarod test at 32 rpm, where hG2019S mice were significantly impaired. Instead, while spontaneous activity did not differ between genotypes, we noticed an impairment in the motor performance at the balance beam and pole test at 6 months, in the rotarod (12 rpm) at 12 and 18 months, in the rotarod (32 rpm) at 6 and 18 months and in the rotarod resistance at 6 and 12 rpm. Finally, 6, 12, and 18 months old hG2019S mice scored significantly worse at the novel object recognition test in terms of mean discrimination index. These observations suggest that 6-months old hG2019S male mice show motor and cognitive impairment.

**Figure 1.**
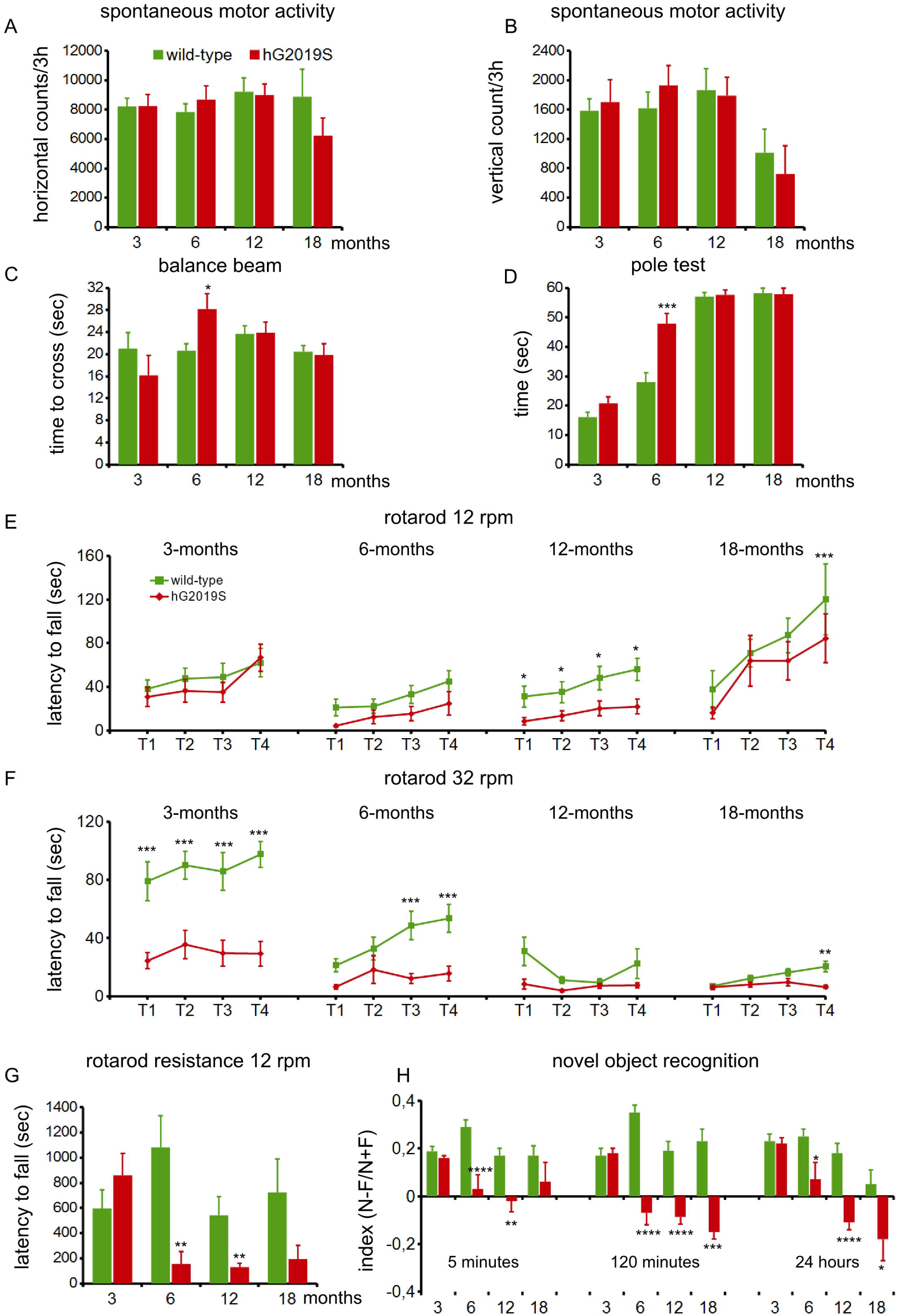
hG2019S mice show age-dependent motor and cognitive impairment. Wild-type and hG2019S mice were profiled for motor and cognitive abilities at 3, 6, 12, and 18 months. 6 and 12 months-old hG2019S mice are characterized by impaired motor coordination and cognitive performance. In detail, we measured spontaneous motor activity in terms of the number of horizontal (A) and vertical counts in 3 hours, time to cross a 6-mm width beam (C), time to reach the ground from a supra-elevated platform (D), time spent on rotarod running at 12 rpm (E) or 32 rpm (F), total resistance on 12 rpm running rotarod (G), and ability to recognize novel object compared to the familiar one (H). Data are shown as mean ±SEM; n=7-18. * p<0.05, ** p<0.01, *** p<0.001,****p<0.0001 versus aged-matched wild-type.

### G2019S mutation correlates with the deposition of NSF aggregates

Protein aggregation is an established histological hallmark of PD. LC3 decorates LB in PD and DLB patients (Higashi et al. 2011; Tanji et al. 2011). Therefore we investigated the presence of proteinaceous aggregates in specimens from substantia nigra, striatum, cortex, and hippocampus obtained from 6-month old wild-type and hG2019S male mice via staining with LC3 antibody. Interestingly, we noticed the presence of a peculiar LC3 staining pattern in nigra, cortex, and hippocampus of hG2091S mice suggesting the presence of perinuclear pale bodies (supplementary figure 1). It has been suggested that LRRK2 mutations promote α-synuclein aggregation (Cresto et al. 2019; MacIsaac et al. 2020). Therefore we studied the distribution of α-synuclein in different brain areas of 6-month old wild-type and hG2019S mice. However, we did not report any overt α-synuclein deposition in the hG2019S mice brain (supplementary figure 2A). LB (Xia et al. 2008) and the related intranuclear inclusion bodies (Pountney et al., 2008) contain more than 90 molecules, including NSF, which we described as LRRK2 functional partner (Piccoli et al. 2014; 2011). Consequently, we first analyzed the subcellular distribution of NSF in hG2019S cortical neurons. We noticed that NSF localizes in somatic structures positive for LC3 and p62 (figure 2A-B), i.e., resembling protein aggregates (Kopito 2000). Protein aggregates form high molecular weight (HMW) complexes that are retained by acetate cellulose in the filter retardation assay (Reid et al. 2003). We confirmed the presence of NSF HMW complex in the sample prepared from 6-months hG2019S mice brain (figure 2C-D) despite a similar expression of LRRK2 and NSF in the two mice lines (supplementary figure 3A-D). A well-established feature of protein aggregates is the resistance to proteinase-K digestion (Kheterpal et al. 2001). Thus we characterized NSF aggregates by limited proteolysis in 6-months old wild-type and hG2019S male mice brain homogenates. Upon proteinase-K digestion, the 100 kDa MW band, corresponding to full-length NSF protein, gradually disappeared in samples prepared from wild-type but not from hG2019S mice (figure 2E-F).

**Figure 2.**
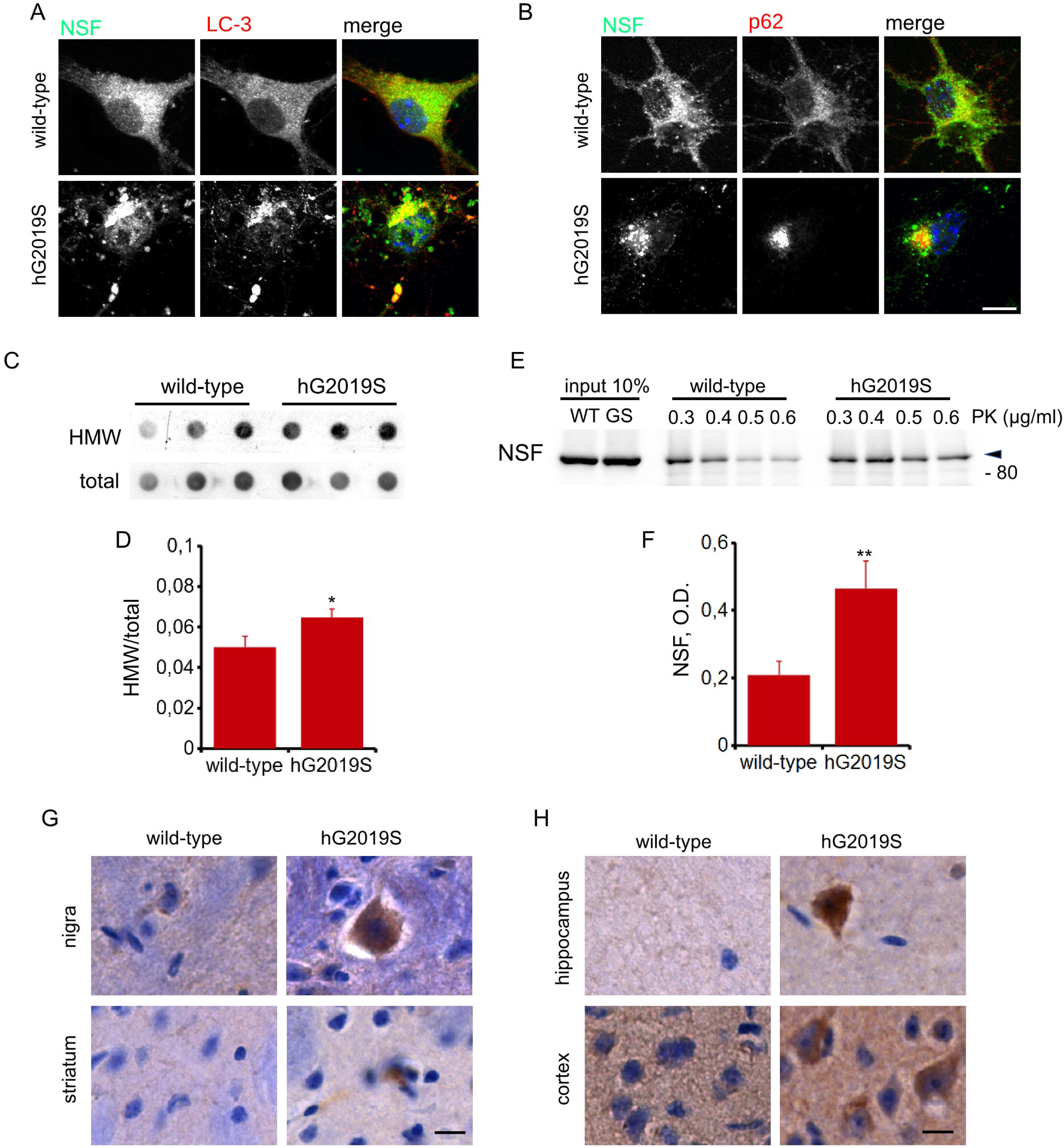
NSF aggregation in hG2019S mice. NSF accumulates in perinuclear aggregates decorated by LC-3 (A) and p62 (B) in DIV14 cortical neurons prepared from hG2019S mice. Images are shown in pseudocolor (NSG green, LC-3, and p62 red, DAPI blue). Scale bar= 10μm. DIV14 cortical cultures from wild-type or hG2019S mice were assayed by filter retardation assay to isolate the high-molecular-weight form of NSF (HMW) or by dot-blot to measure total NSF protein (C). The graphs report NSF aggregation expressed as HMW fold-over total NSF. Data are shown as mean ±SEM, n=8. * p<0.05 versus wild-type (D). The biochemical analysis of brain homogenate shows that NSF is resistant to proteinase K degradation in samples prepared from of 6-months old hG2019S mice. Arrowhead indicates the band corresponding to full-length NSF (E). The graphs report the amount of full-length NSF protein expressed as optical density after digestion with 0.5 μg/ml proteinase K. Data are shown as mean ±SEM, n=8. * p<0.05 versus wild-type (F). NSF accumulates in aggregates in substantia nigra and striatum (G) as well as in cortex and hippocampus (H) in 6-months hG2019S LRRK2 mice. Scale bar= 10μm.

Histological analysis revealed the presence of NSF positive aggregates in nigral, striatal, hippocampal, and cortical sections prepared from 6-months hG2019S mice (figure 2G-H). Next, we characterized post-mortem basal ganglia specimens obtained from healthy control, idiopathic, and G2019S PD cases. The detergent solubility profile characterizes proteinaceous aggregates from the biochemical standpoint (Mamais et al. 2013). Interestingly, while Triton-X100 soluble NSF levels were comparable among the groups (figure 3A, C), the yield of NSF in the SDS soluble fraction was higher in the sample prepared from idiopathic or G2019S PD patients (figure 3B, D). Besides, we found that NSF decorates a proportion of α-synuclein positive structure in both idiopathic and G2019S PD cases (figure 3E and table 1). To directly assess the impact of G2019S mutation on the NSF aggregation state, we analyzed dopaminergic neurons differentiated from G2019S human neuronal precursor and their respective gene-corrected line (supplementary figure 3E-F). By filter retardation assay, we noticed that NSF forms HMW complexes in samples prepared in G2019S dopaminergic neurons (figure 3F-G). Altogether this evidence suggests that LRRK2 G2019S mutation correlates with the deposition of NSF aggregates in both murine and human models.

**Figure 3.**
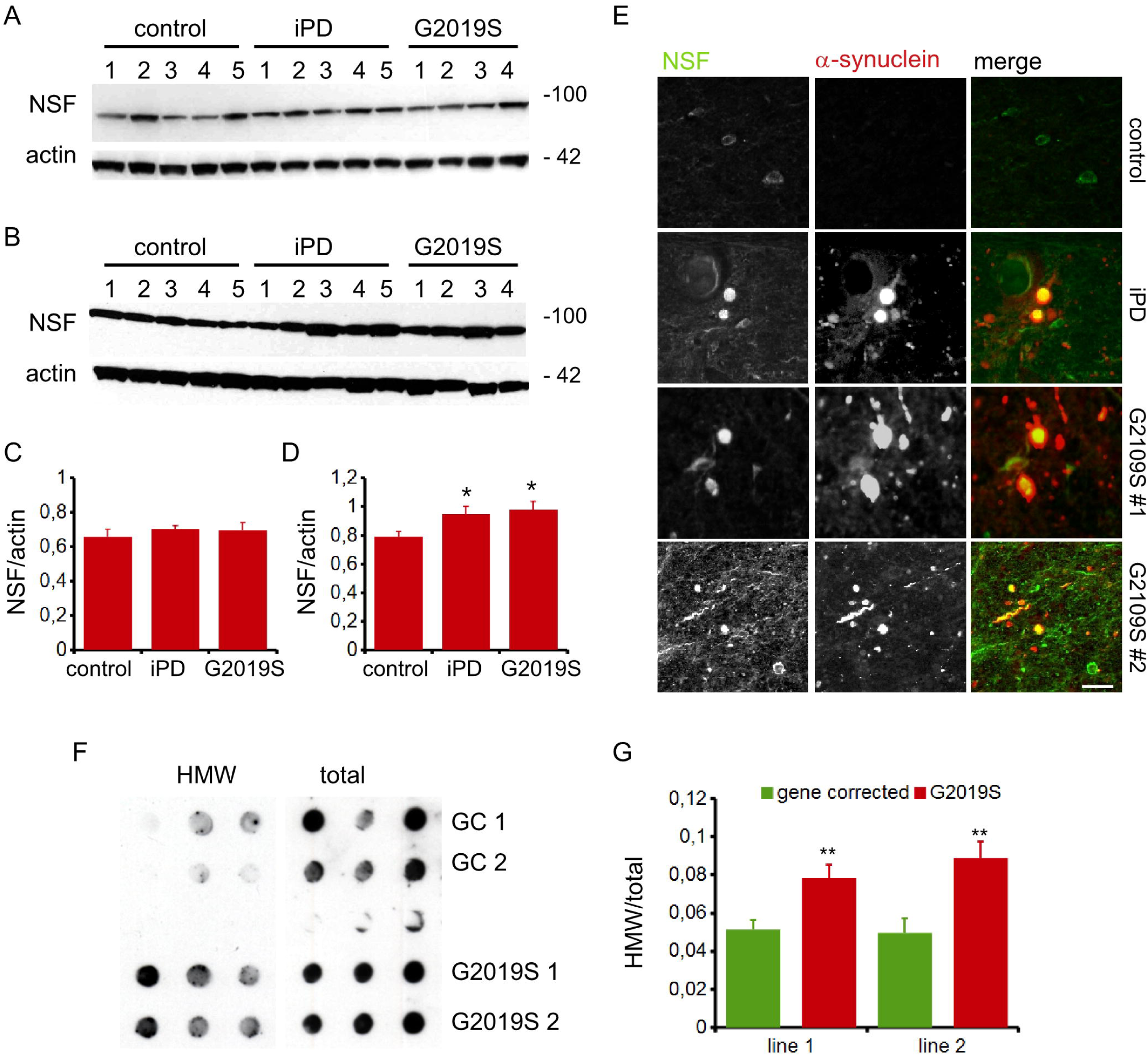
NSF in human-derived samples. NSF distribution in the Triton-X100 (A) or SDS (B) soluble fraction in PD specimens. Graph reports NSF optical density in the Triton-X100 (C) or SDS (D) soluble fraction, normalized versus actin amount. Data are shown as mean ±SEM, n=4-5. * p<0.05 versus control. Post-mortem analysis of nigra specimens shows that NSF decorates α-synuclein positive Lewy bodies and Lewy neurite in G2019S PD patients. Scale bar= 10μm (E). Induced dopaminergic neurons differentiated from two independent G2019S patients (G2019S 1 and 2) as well as gene-corrected counterpart (GC 1 and 2) were assayed by filter retardation assay to isolate high-molecular-weight form of NSF (HMW) or by dot-blot to measure total NSF protein (total) (F). The graphs report NSF aggregation expressed as HMW fold-over total NSF. Data are shown as mean ±SEM, n=8. ** p<0.001 versus wild-type (G).

**Table 1.**
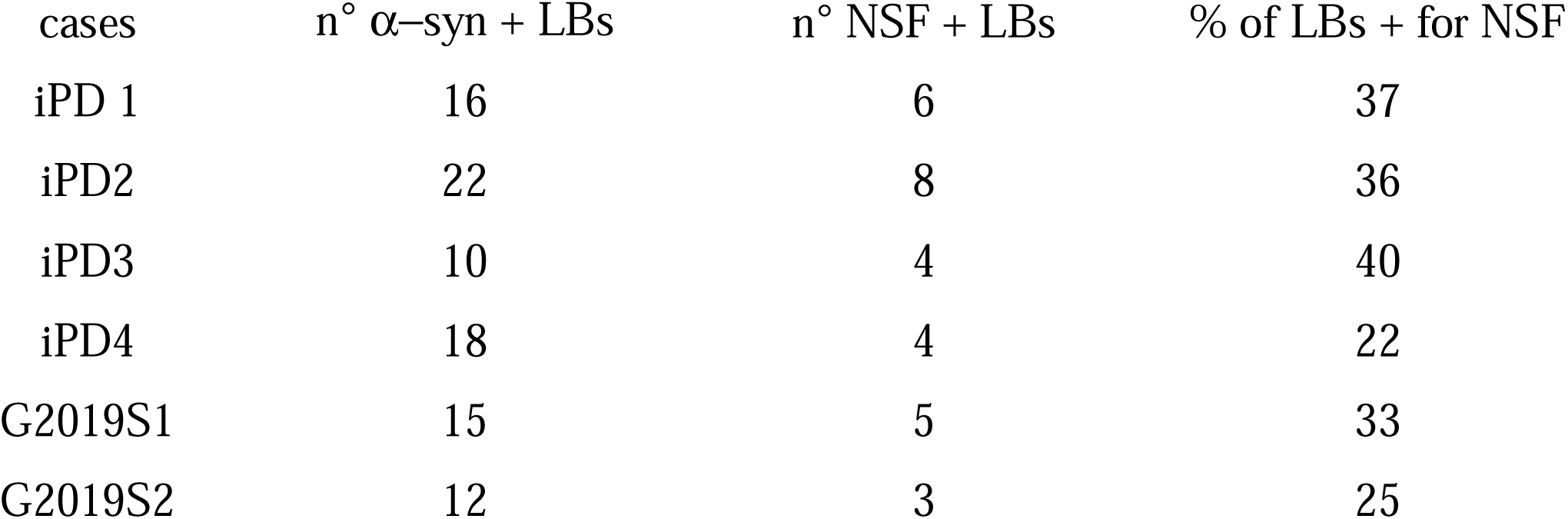
The table lists the number of α-synuclein positive structure (α-syn +), the number of α-synuclein and NSF positive structure (NSF +) and the relative percentage (%) in specimens prepared from substantia nigra region of 4 idiopathic (iPD) and 2 G2019S patients

| cases | n° α-syn + LBs | n° NSF + LBs | % of LBs + for NSF |
| --- | --- | --- | --- |
| iPD 1 | 16 | 6 | 37 |
| iPD2 | 22 | 8 | 36 |
| iPD3 | 10 | 4 | 40 |
| iPD4 | 18 | 4 | 22 |
| G2019S1 | 15 | 5 | 33 |
| G2019S2 | 12 | 3 | 25 |

### LRRK2 G2019S kinase activity triggers NSF aggregation

Previously we have shown that NSF is an LRRK2 substrate *in vitro* (Belluzzi et al. 2016). To complement this finding, we generated an antibody specifically recognizing NSF phosphorylated at Thr645 (supplementary figure 4A-B). This reagent showed that NSF phosphorylation at Thr645 increased in post-mortem samples gathered from G2019S patients specimen (figure 4A-C) and in several brain areas gathered from 6-months hG2019S mice (figure 4D-E). Structural analysis of NSF (PDB ID: 3J94) revealed that the side chain of T645 is close to the ATP-binding site on the D2 domain involved in NSF oligomerization (aa 490-744) (figure 4F-G). By differential detergent solubility assay, we noticed an increased amount of NSF in the insoluble fraction gathered from hG2019S cortical neurons (figure 4H-I). We obtained a similar outcome upon the overexpression of hLRRK2 G2019S in N2A cell (supplementary figure 4C-D). This biochemical evidence suggests that LRRK2 G2019S triggers NSF aggregation. Proteins can aggregate upon impairment of proteasome activity (Varshavsky, 2017). Accordingly, a filter retardation assay showed that the pharmacological proteasome inhibition induced NSF aggregation in wild-type neurons. Instead, hG2019S neurons were characterized by a high basal amount of HMW NSF that did not further increase upon MG-132 treatment (supplementary figure 4E-F). Indeed, LRRK2 G2019S might interfere with UPS mediated NSF degradation. However, we reported a comparable UPS proteolytic activity in 6- or 12-months old wild-type and hG2019S mice (supplementary figure 4G). Therefore, we thought that other mechanisms must underline LRRK2 G2019S dependent NSF aggregation. Phosphorylation may directly affect protein folding and oligomeric state (Metskas e Rhoades 2015). To dissect the biological consequence of NSF phosphorylation on Thr645, we generated by site-direct mutagenesis the NSF phosphomimetic variant NSF T645D. As expected (Belluzzi et al. 2016), NSF T645D has an augmented ATPase activity (supplementary figure 5A-B). NSF T645D is prone to aggregation independently from UPS blockade (supplementary figure 5C-D) and it forms higher-molecular weight complexes than the wild-type variant (supplementary figure 5E-F). NSF T645D is strongly enriched in the insoluble fraction, as suggested by differential detergent solubility (figure 4J-K). Strikingly, once co-expressed with HA-NSF wild-type, NSF T645D influenced the solubility of the wild-type protein (figure 4J, L). These observations suggest that phosphorylation at T645 triggers NSF aggregation. Once entrapped within aggregates, proteins get cleaved and tagged by ubiquitin (Olanow et al. 2004). Our western-blotting analysis revealed that NSF T645D undergoes proteolytic cleavage and is strongly decorated by ubiquitin (figure 4M-O). In ubiquitin chains, each monomer can be conjugated through different lysine residues, generating a code with functional consequences [reviewed in (Akutsu, Dikic, e Bremm 2016)]. To determine the nature of ubiquitination occurring on the two NSF variants, we expressed in HEK293 cells NSF WT and NSF T645D together with ubiquitin wild-type or with individual ubiquitin constructs missing lysine 48, 48 and 63 or 48, 63 and 29, respectively. We noticed that the degradation and the ubiquitination of the two NSF variants depend on different lysine residues (supplementary figure 5G-K). Altogether, these data suggest that LRRK2 phosphorylation at T645 alters NSF biochemical properties and promotes its aggregation.

**Figure 4.**
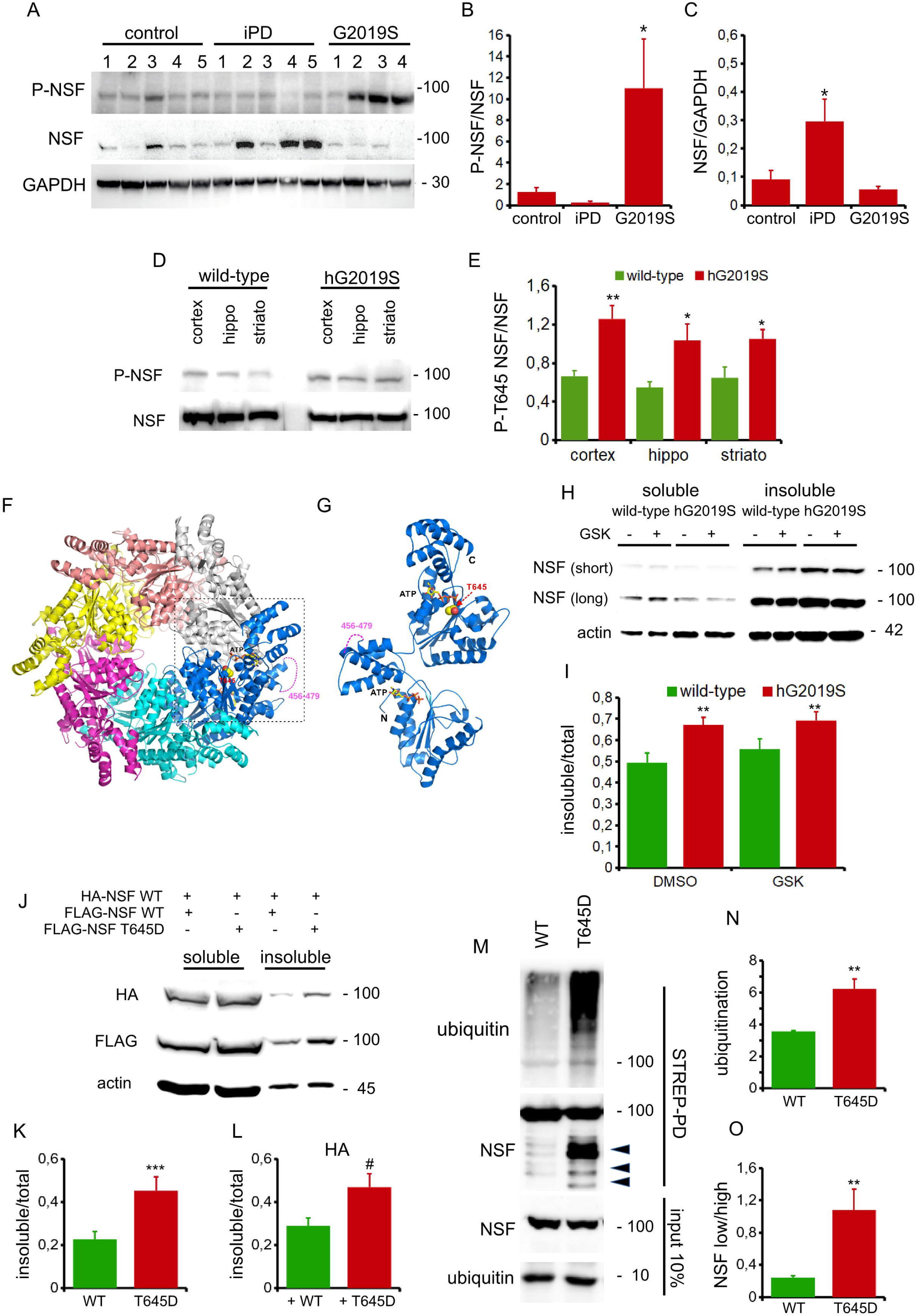
LRRK2 phosphorylation induces NSF aggregation. NSF is phosphorylated at Thr645 in G2019S PD-patients brains (A). The graphs report NSF phosphorylation at Thr645 fold-over total NSF (B) as well as NSF relative amount (C). Data are shown as mean ±SEM, n=8. * p<0.05. NSF is phosphorylated at Thr645 in hG2019S mice brain (D). The graphs report NSF phosphorylation at Thr645 fold-over total NSF (E). Data are shown as mean ±SEM, n=8. *, ** p<0.05, 0.01 versus wild-type. The combined ribbon, stick, and sphere representation showing the overall architecture of NSF hexamer structure from human (PDB ID: 3J94) as well as the positions of T645 and two bound ATP molecules in one NSF monomer. In this drawing, the six NSF monomers are shown in different colors, and the side chain of T645 is shown in the sphere mode and the bound ATP in the stick mode. The missing region between residue 456 and 479 in the NSF monomer is represented by a dashed pink line (F). The enlarged view of an NSF monomer in blue in panel G. Notably, the side chain of T645 is very close to the bound ATP molecule on the D2 domain (G). LRRK2 phosphorylation influences NSF aggregation. We assayed the solubility profile of NSF in samples prepared from DIV14 wild-type or hG2019S cortical neurons and treated with vehicle (DMSO) or the LRRK2 kinase inhibitor GSK2578215A (GSK, 0.2 μM, 18 hours). Short and long acquisition time of the ECL signal emitted by anti-NSF western-blot is reported (H). The graphs report the amount of NSF present in the Triton-X100 insoluble fraction, expressed as fold-over total NSF. Data are shown as mean ±SEM, n=11. ** p<0.01 versus wild-type DMSO (I). We assayed the solubility profile of FLAG- and HA-NSF in samples prepared from HEK293 cell over-expressing wild-type or phosphomimetic T645D Strep-FLAG-NSF isoform together with HA-NSF wild-type. Strep-FLAG NSF T465D is present in the Triton-X100 insoluble fraction and sequesters HA-NSF wild-type (J). The graphs report the amount of FLAG-NSF (K) and HA-NSF (L) present in the Triton-X100 insoluble fraction, expressed as fold-over total NSF. We transfected in HEK293T cells myc-ubiquitin together with Strep-FLAG-NSF WT or T645D. We purified on streptavidin-beads NSF protein and assessed ubiquitination level by western-blotting with anti-myc antibody. Arrowheads indicate putative NSF low molecular weight fragments detected by anti-FLAG antibody (M). The graphs report NSF ubiquitination in presence of myc-ubiquitin wild-type, calculated as anti-myc optical density normalized versus total NSF yield (N) and NSF fragmentation, expressed as the ratio of low molecular weight NSF fragments over high molecular weight NSF optical density (O). Data are shown as mean ±SEM, n=7. ** p<0.01 versus wild-type.

### NSF over-expression is detrimental for hG2019S neurons in an LRRK2 kinase-dependent manner

NSF plays a crucial role in neurons (Bonnycastle, Davenport, e Cousin 2020). To investigate whether NSF aggregation may be neurotoxic, we infected wild-type and hG2019S neurons with lentiviral viruses expressing mCherry alone or together with NSF wild-type, the phospho-null variant NSF T645A or the phospho-mimetic variant NSF T645D. Neurons were treated from DIV10 to DIV14 with DMSO or GSK2578215A (GSK, 0.2 μM daily) and then processed at DIV16 (figure 5A and supplementary figure 6A). Neurite retraction is an established marker of neuronal damage (Chernova et al. 2007). Our morphological analyses revealed that the over-expression of either NSF wild-type or T645A had no major effect on wild-type neuron morphology. Instead, NSF T645D was detrimental to wild-type neurons. Noteworthy, NSF wild-type over-expression caused a significant reduction of neurite number in hG2019S neurons. Such morphological phenotype was absent in hG2019S neurons over-expressing the phospho-null NSF T645A variant (figure 5B). The analysis of neuronal viability via MTT assay confirmed this outcome (figure 5C). Interestingly, the chronic treatment with the LRRK2 kinase inhibitor GSK2578215A prevented the neurite loss of hG2019S cultures expressing wild-type NSF but did not ameliorate the overall viability. Cumulatively, these data indicate that NSF is neurotoxic upon LRRK2 phosphorylation.

**Figure 5.**
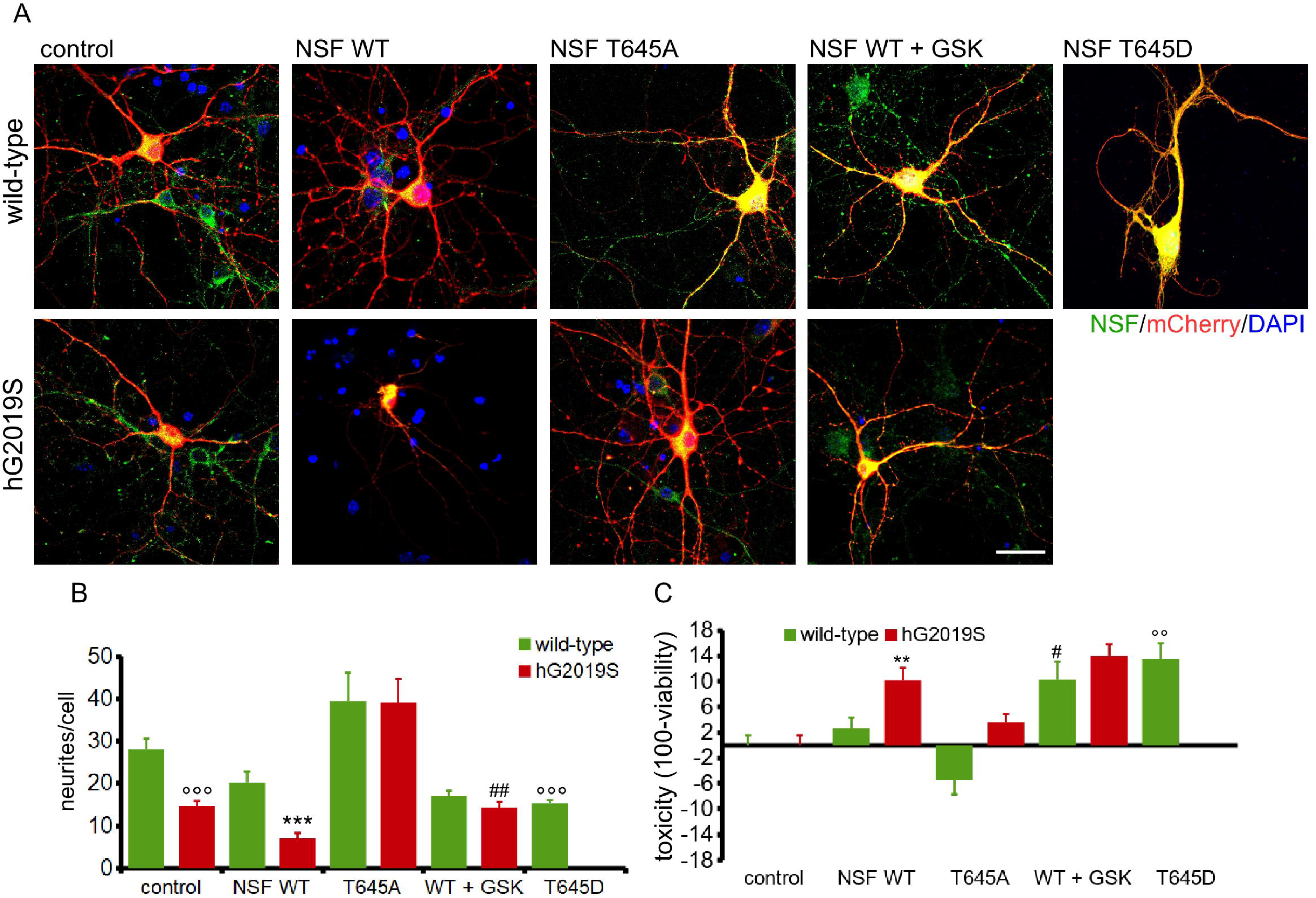
LRRK2 phosphorylation triggers NSF toxicity. Representative images of DIV14 Wild-type and hG2019S cortical neurons transduced at DIV4 with mCherry expressing viruses (control) or viruses co-expressing mCherry and wild-type Strep-FLAG NSF WT, phospho-null variant T645A or T645D. Where indicated, cells were chronically treated from DIV4 to DIV14 with DMSO or GSK2578215A (GSK, 0.2 μM, every two days). Cells were then processed for imaging purposes at DIV14, scale bar= 40um (A). The graph reports the number of neurites per neuron. Data are shown as mean ±SEM, n=14. *** p<0.001 versus NSF WT in wild-type neuron, °°° p<0.001 versus control infection in wild-type neuron, ## p<0.01 versus NSF WT in hG2019S neurons DMSO (B). Neuronal cultures were infected and treated as above and then processed for MTT assay at DIV14. The graph reports toxicity, expressed as 100-relative viability (C). Data are shown as mean ±SEM, n=18. ** p<0.01 versus NSF WT in wild-type neurons, # p<0.01 versus NSF WT in wild-type neurons DMSO,°° p<0.001 versus control infection in wild-type neuron.

### Trehalose treatment clears NSF aggregates

Autophagy can attenuate protein aggregation (Fujikake, Shin, e Shimizu 2018). We used trehalose, a well-established autophagic inducer (DeBosch et al. 2016), to stimulate autophagy in wild-type and hG2019S neurons (supplementary figure 6B-C). The filter retardation assay suggested that trehalose treatment reduced NSF aggregation in NSF over-expressing HEK293 cell treated with MG132 (supplementary figure 6D-E). *In vivo*, trehalose treatment reduced NSF aggregation in 6-months old hG2019S male mice (1% in the drinking water, 1-month; supplementary figure 6F-G) without significant impact on animal weight (supplementary figure 6H). Specifically, trehalose treatment reduced the number of cells decorated by NSF in substantia nigra (figure 6), striatum (figure 7), cortex, and hippocampus (figure 8) region of hG2019S mice. Caspase-3 cleavage precedes apoptotic cell death. We noticed a significant increase of cleaved caspase-3 positive cells in nigra (figure 6) and striatum (figure 7) without any overt loss of TH density. Caspase-3 positive cells were also evident in the cortex and hippocampus (figure 8) of hG2019S mice. Trehalose treatment significantly reduced the number of caspase-3 positive cells in substantia nigra (figure 6), striatum (figure 7), cortex, and hippocampus (figure 8) of hG2019S mice. Our observations suggest that trehalose treatment cleared NSF aggregation and reduced cell death in hG2019S mice brain.

**Figure 6.**
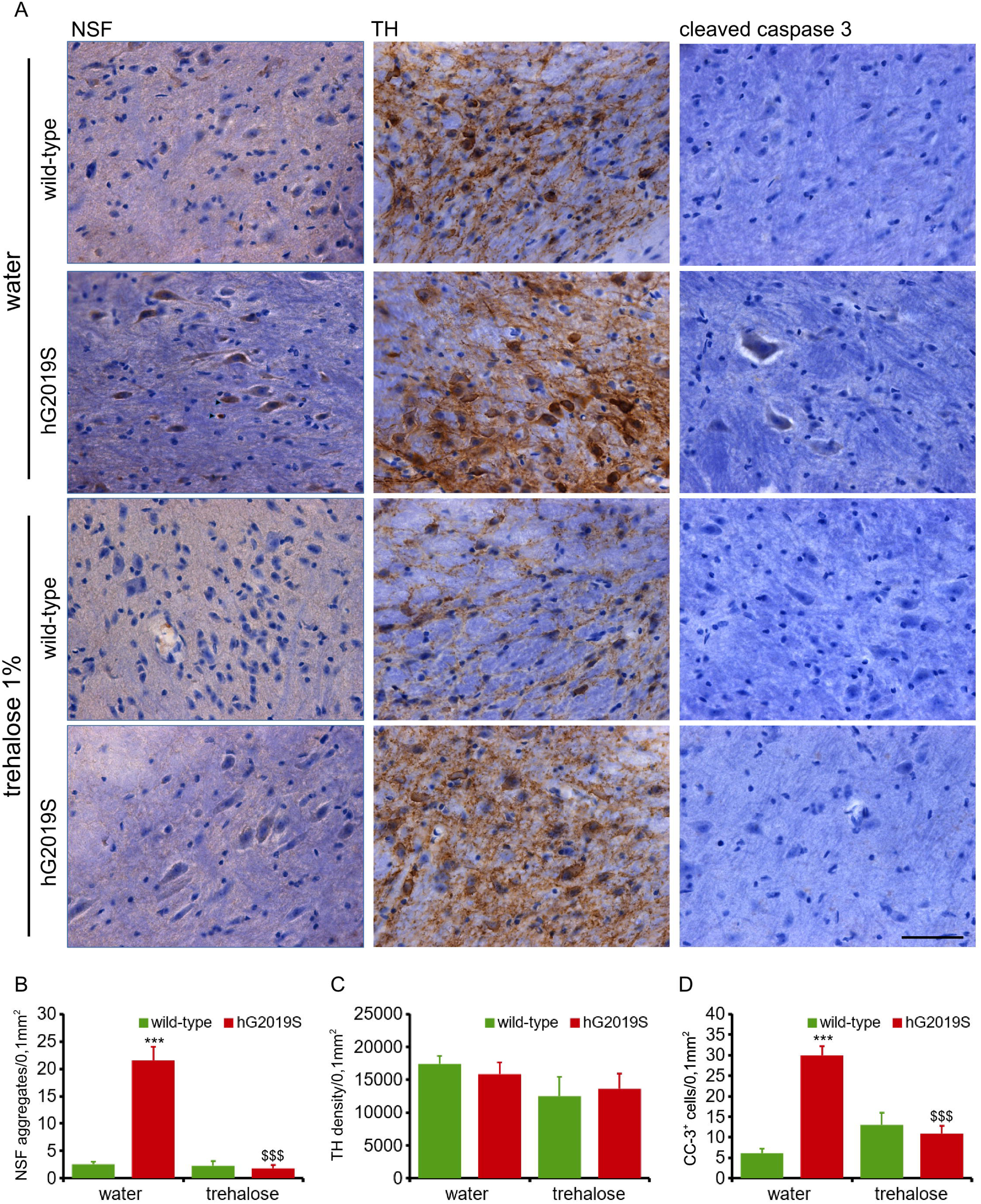
Trehalose treatment rescues histochemical pathological hallmark of hG2019S mice in the substantia nigra. Wild-type and hG2019S mice were treated starting at 5 months with trehalose (1% in drinking water) and processed for imaging analysis at 6 months. Nigra sections were stained with anti-NSF, anti-TH, and anti-cleaved caspase-3 antibodies and counterstained with hematoxylin to visualize nuclei. hG2019S mice are characterized by NSF aggregation and caspase-3 cleavage in nigra. Trehalose treatment reduces NSF aggregation and cleaved-caspase 3 signal, scale bar = 50 μm (A). The graph reports the number of NSF aggregates (B), the TH optical density (C), and the number of caspase-3 positive cells (D) detected in a 0.1 mm^2^ area. Data are shown as mean ±S.E, n=6. *** p<0.001 versus wild-type, same treatment, $$$ p<0.001 versus water, same genotype.

**Figure 7.**
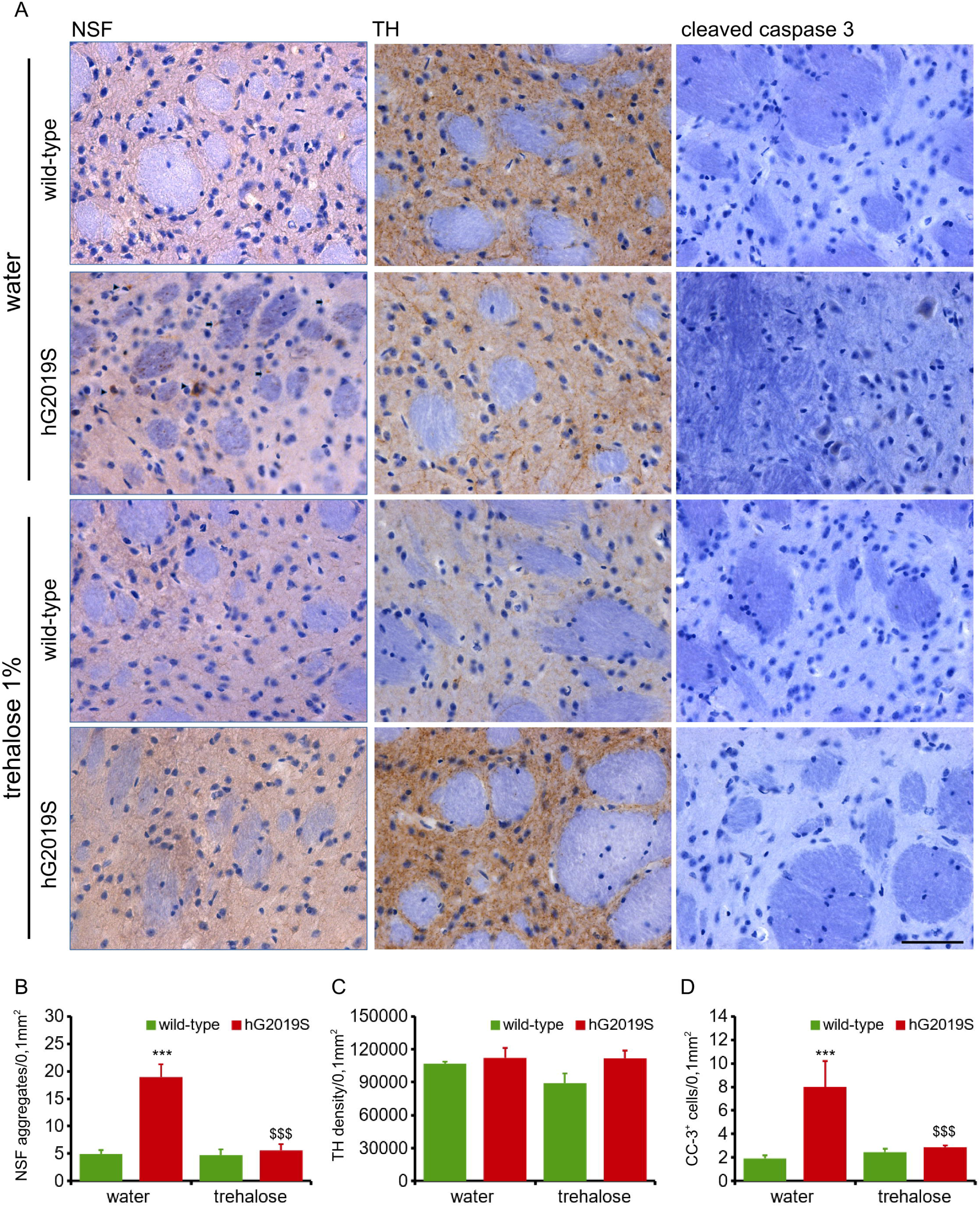
Trehalose treatment rescues the histochemical pathological hallmark of hG2019S mice in the striatum. Wild-type and hG2019S mice were treated starting at 5 months with trehalose (1% in drinking water) and processed for imaging analysis at 6 months. Striatal sections were stained with anti-NSF, anti-TH, and anti-cleaved caspase-3 antibodies and counterstained with hematoxylin to visualize nuclei (A). The graph reports the number of NSF aggregates (B), the TH optical density (C) and the number of caspase-3 positive cells (D) detected in a 0.1 mm^2^ area. Data are shown as mean ±S.E, n=6. *** p<0.001 versus wild-type, same treatment, $$$ p<0.001 versus water, same genotype.

**Figure 8.**
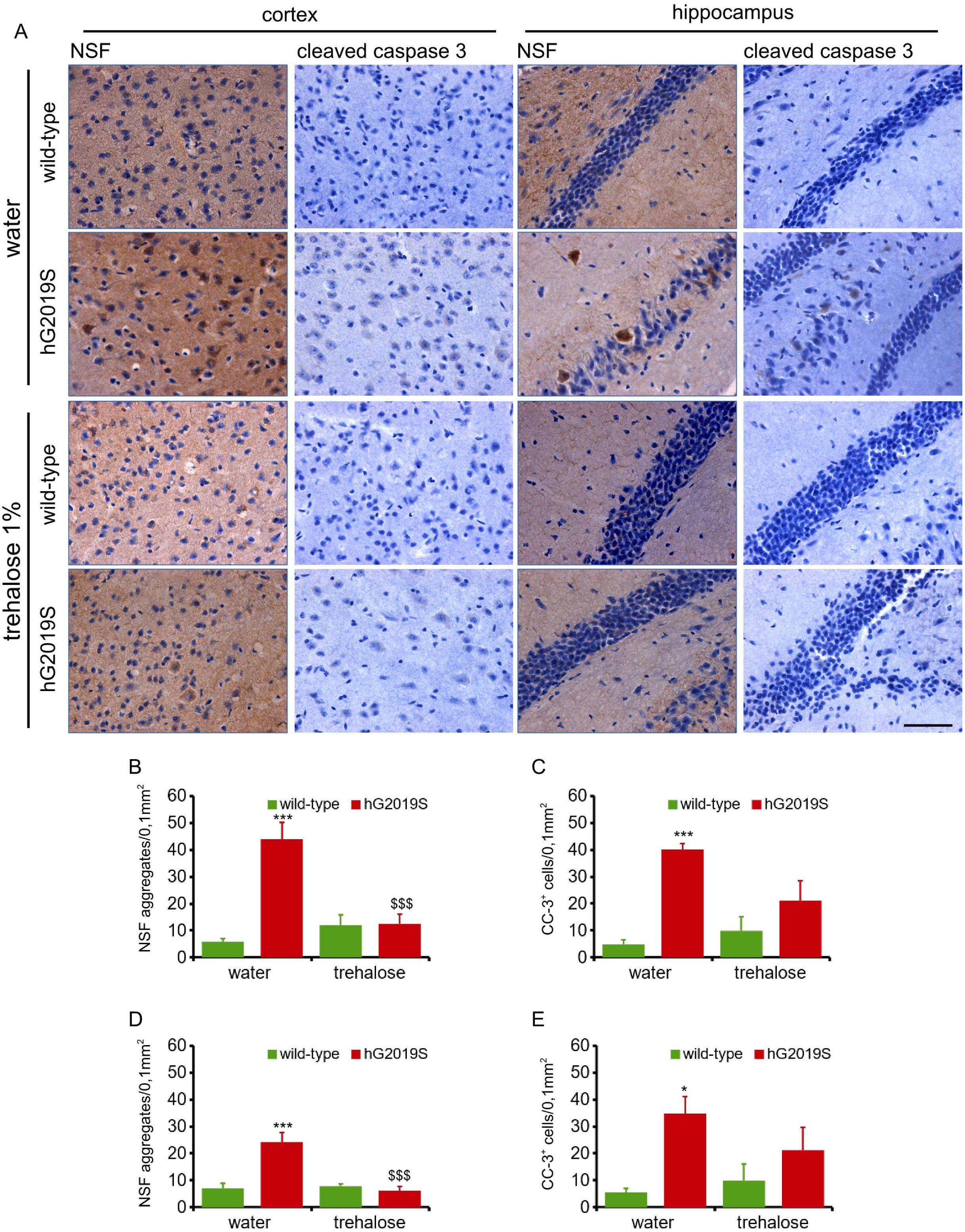
Trehalose treatment rescues the histochemical pathological hallmark of hG2019S mice in the cortex and hippocampus. Wild-type and hG2019S mice were treated starting at 5 months with trehalose (1% in drinking water) and processed for imaging analysis at 6 months. Brain sections encompassing cortical area or hippocampus were stained with anti-NSF and anti-cleaved caspase-3 antibodies and counterstained with hematoxylin to visualize nuclei. hG2019S mice are characterized by NSF aggregation and caspase-3 cleavage in the cortex and hippocampus. Trehalose treatment reduces NSF aggregation and cleaved-caspase 3 signal (A). The graph reports number of NSF aggregates in cortex (B) and hippocampus (C) as well as the number of caspase-3 positive cells in cortex (D) and hippocampus (E) detected in a 0.1 mm^2^ area. Data are shown as mean ±S.E, n=5. *, *** p<0.05, p<0.001 versus wild-type, same treatment, $$$ p<0.001 versus water, same genotype.

### Trehalose treatment ameliorates motor and cognitive phenotype in hG2019S mice

Next, we profiled the motor and cognitive performances in 6-month old hG2019S and wild-type male mice upon chronic treatment with trehalose (figure 9). We reported a robust recovery of motor and cognitive abilities in trehalose-treated animals. In particular, trehalose treatment ameliorated the performances at the 6-mm balance beam (figure 9C), at the 32-rpm rotarod (figure 9F), the rotarod resistance at 12 rpm (figure 9G) and the novel object recognition (figure 9H). We also observed a partial improvement in motor and cognitive performances (in the rotarod 32 rpm and in the novel object recognition) in 12-month old hG2019S mice treated with trehalose starting at month 10 (1% in the drinking water, 2-month; supplementary figure 7). Altogether, these observations suggest that trehalose treatment is beneficial in rescuing behavioral defects of hG2019S mice.

**Figure 9.**
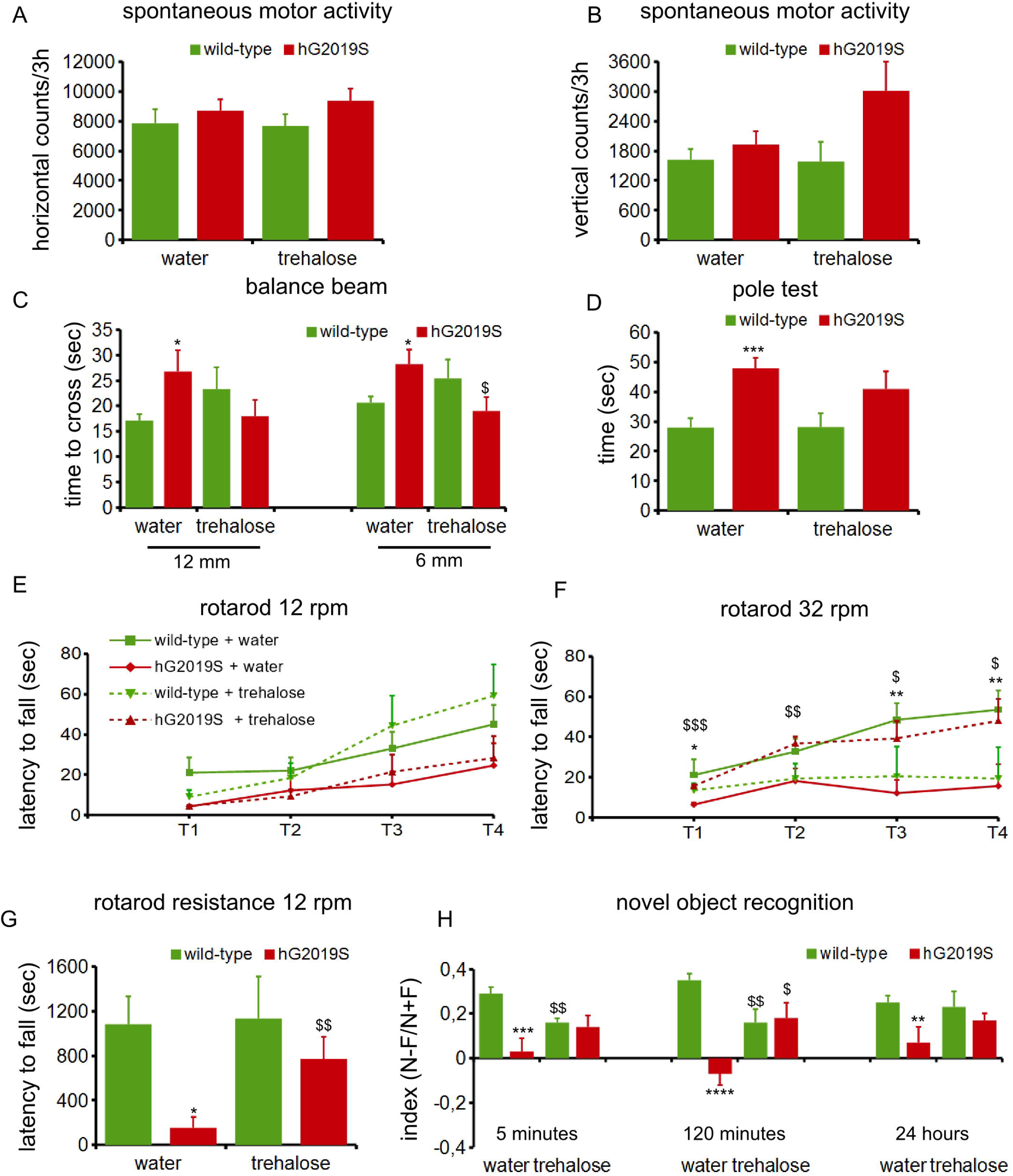
Trehalose treatment rescues motor and cognitive defects of hG2019S mice. Wild-type and hG2019S mice were treated starting at 5 months with trehalose (1% in drinking water) and profiled for motor and cognitive abilities at 6 months. hG2019S mice are characterized by impaired motor coordination and cognitive performance. In detail, we measured spontaneous motor activity in terms of the number of horizontal (A) and vertical (B) counts in 3 hours, time to cross a 12- and 6-mm width beam (C), time to reach the ground from a supra-elevated platform (D), time spent on rotarod running at 12 rpm (E) or 32 rpm (F) over 4 trials (T_1_-T_4_), total resistance on 12 rpm running rotarod (G), and ability to recognize novel object compared to the familiar one (H). Data are shown as mean ±SEM; n=7-18. * p<0.05, ** p<0.01, *** p<0.001, **** p<0.0001 versus wild-type, same treatment, $ p<0.05, $$ p<0.01, $$$ p<0.001 versus water, same genotype.

## Discussion

### LRRK2 and the proteostatic control

In the present study, we show that hG2019S mice present an age-dependent motor and cognitive impairment together with the deposition of protein aggregates containing NSF and signs of cell death in substantia nigra, striatum, cortex, and hippocampus. Post-mortem brain investigation demonstrated that LRRK2 G2019S patients often show synucleinopathy, occasionally tauopathy, suggesting a role for LRRK2 in protein inclusion pathology (Taymans e Cookson 2010). The description of frontotemporal ubiquitinated neuronal intranuclear inclusions in one patient carrying G2019S substitution indicates that the LRRK2 pathological impact is not restricted to the substantia nigra (Dächsel et al. 2007; Kalia et al. 2015).

LRRK2 kinase activity brings to protein accumulation without affecting the catalytic activity of the proteasome or expression levels of proteasomal core subcomplexes (Lichtenberg et al. 2011). Instead, LRRK2 influences the autophagic pathways at multiple levels [reviewed in (Manzoni e Lewis 2017; Albanese, Novello, e Morari 2019; Cogo et al. 2020). The investigation of *LRRK2*-deficient mice revealed a biphasic alteration of autophagy in the kidney (Tong et al. 2010a; 2012); however, similar studies in an independent *LRRK2*-knock out mouse line demonstrated an accumulation of secondary lysosomes in the kidney without major involvement of autophagy (Herzig et al. 2011a). Even more complicated is the state-of-art describing autophagy in an LRRK2 disease context. Many authors conclude that G2019S mutation increases autophagy in different models, including stable cell lines (Gómez-Suaga et al. 2012; Plowey et al. 2008), fibroblasts (Yakhine-Diop et al., 2014), iPS derived neurons (Bravo-San Pedro et al. 2013), *C*.*Elegans* (Ferree et al., 2012) and mice (Ramonet et al., 2011). However, other studies have reported that the same mutation reduces (Manzoni et al. 2013) or at least does not affect (Sánchez-Danés et al. 2012; Wauters et al. 2019) autophagy. Such conflicting results leave unclear the impact of LRRK2 G2019S mutation on autophagy (Manzoni 2017). Still, increased accumulation of protein aggregates has been reported in complementary LRRK2 G2019S models (Bang et al. 2016; Guerreiro et al. 2016; Novello et al. 2018; Schapansky et al. 2018). Thus it might be possible that LRRK2 controls protein clearance acting upstream of the two main protein-clearance mechanisms.

### Phosphorylation can trigger protein aggregation

Post-translational modifications regulate protein structure. In particular, phosphorylation can dictate protein folding and eventually, aggregation state. In the context of PD, phosphorylation has a severe impact on alpha-synuclein [reviewed in (Tenreiro, Eckermann, e Outeiro 2014)]. While soluble, monomeric alpha-synuclein is largely not phosphorylated in physiological conditions, up to 90% of LB-synuclein is phosphorylated on Ser-129 (Anderson et al. 2006). Indeed it is still unresolved whether phosphorylation might trigger or impair alpha-synuclein aggregation or toxicity. While several reports claim that phosphorylation at Ser-129 favors protein aggregation (Kragh et al. 2009; Wu et al. 2011) other authors described opposite or no effect (Fiske et al. 2011; Sancenon et al. 2012). However, phosphorylation at Ser-129 influences alpha-synuclein sub-cellular localization (Gonçalves e Outeiro 2013), suggesting that such PTM dictates alpha-synuclein function and fate. Similarly, hyperphosphorylated Tau protein precipitates in toxic aggregates and destabilizes the axonal tubulin cytoskeleton (Alonso et al. 2018). In vitro evidence shows that the phosphorylation of specific Tau residues modifies local folding, thus affecting global structure (Bibow et al. 2011; Inoue et al. 2012). We showed that LRRK2 phosphorylates NSF on Thr-645 *in vitro* (Belluzzi et al. 2016) as well as *in vivo* (here). Based on our structural analysis, the side chain of T645 falls in proximity to the ATP-binding site on NSF located on the D2 domain (aa 206-477). In particular, Thr-645 lies very close to the phosphate group of the bound ATP molecule. ATP binding on the D2 domain is essential for the formation of NSF hexamer (Whiteheart, Schraw, e Matveeva 2001). We determined by complementary means that upon phosphorylation at Thr-645 NSF forms *bona fide* HMW aggregates, being insoluble to Triton-X100 (Naslavsky et al. 1997), resistant to proteinase-K treatment (Kheterpal et al., 2001) and decorated by ubiquitin and p62 (Bjørkøy et al. 2005). Altogether, our data suggest that the phosphorylation of NSF at T645 promotes NSF oligomerization and eventually triggers its precipitation into protein inclusions. Indeed it is unlikely that LRRK2-mediated phosphorylation is sufficient by itself to cause NSF aggregation. We reported that Rab3A, a well-established LRRK2 substrate (Lis et al. 2018; Steger et al. 2016), does not aggregate in BAC hG2019S mice (supplementary figure 2B). Thus, LRRK2 phosphorylation might not be the unique driver promoting NSF aggregation. Actually, in idiopathic PD patients, we noticed that NSF is expressed at a high level and decorates a proportion of LB. This evidence may argue against a causative role for LRRK2 kinase activity. However, it has been recently postulated a role for LRRK2 in idiopathic patients (Di Maio et al. 2018). It is tempting to speculate that in presence of increased NSF level even the low kinase activity characterizing wild-type LRRK2 may be sufficient to promote NSF aggregation. As reported for other aggregation-prone proteins like tau, it is conceivable that NSF has an intrinsic propensity towards aggregation that is pathologically exacerbated upon LRRK2 phosphorylation.

### NSF aggregation, aging, and neuronal toxicity

NSF is a key component of presynaptic machinery, allowing the first step of SV recycling (Pallanck et al. 1995). Numerous studies have shown that depletion of cytosolic NSF impair membrane fusion machinery (Rothman, 1994) and results in the accumulation of intracellular vesicles (Malhotra et al. 1988; Mohtashami et al. 2001). Thus, NSF aggregation might affect presynaptic fusion machinery. Interestingly, it has been reported that experimental ischemia induces NSF aggregation into Triton-X100 insoluble inclusions that harm neuronal function via sequestering synaptic vesicle (Liu e Hu 2004). A presynaptic dysfunction might indeed explain the early-stage manifestation of PD observed in animal models (Longo et al. 2017; Melrose et al. 2010; Sloan et al. 2016) or presymptomatic LRRK2 G2019S carriers (Sossi et al. 2010). However, it is difficult to envisage how aberrant vesicle release could cause the overt neuronal death that characterizes the late phase of the disease. Neurons benefit from several pathways to effectively handle protein aggregation, but the ultimate resource to counteract proteinaceous stress is the degradative clearance of misfolded protein (Rubinsztein 2006). Nonetheless, the structure of the proteasome complex itself disfavors the removal of large oligomeric and aggregated proteins: substrates need to be unfolded to pass through the tight pore of the proteasome barrel (Verhoef et al. 2002). Autophagy may help, being capable of handling large protein aggregates (Hara et al. 2006; Komatsu et al. 2006). Unfortunately, both autophagy and proteasome activity pronouncedly declines with aging (Del Roso et al. 2003; Graham e Liu 2017; Martinez-Vicente, Sovak, e Cuervo 2005). Thus, aging and pathological LRRK2 phosphorylation together may contribute to the deposition of NSF aggregates. Given the low stoichiometry that we reported for LRRK2-driven NSF phosphorylation *in vitro* (Belluzzi et al. 2016), it may well be that NSF phosphorylation occurs rarely *in vivo*. However, we noticed that T465D NSF affects the detergent solubility of the unmodified protein. Thus, we may depict a model where few phosphorylated NSF molecules lead the formation of pathological seeds capable of sequestering unphosphorylated protein into large aggregates that accumulate along with aging.

### Hints toward a therapy

Altogether this evidence may call for a therapeutic strategy targeting LRRK2 kinase activity. Several brain penetrant LRRK2 selective inhibitors have been identified [reviewed in (Ding e Ren 2020)]; however the broad expression of LRRK2 in other organs apart from the central nervous system, including lung, kidney and the immune system raise issues about side effects (Baptista et al. 2013; Fuji et al. 2015; Herzig et al. 2011b; Ness et al. 2013; Tong et al. 2010b; Baptista et al. 2020). In our model, LRRK2 kinase inhibition resulted partially toxic and not efficacious in resolving NSF aggregation. Thus, complementary therapeutic approaches are still needed. Targeting the protein aggregates caused by LRRK2 kinase activity may be clinically relevant. It is reasonable to predict that LRRK2 kinase activity affects the biochemical profile of other substrates besides NSF. Thus, a more holistic solution is needed. The activation of autophagy may promote the clearance of toxic protein aggregates. We report that chronic treatment with trehalose reduced NSF aggregation and ameliorated motor and cognitive phenotype in aged G2019S mice. Treatment with trehalose already demonstrated to be protective in models of Huntington disease, spinocerebellar ataxia, Machado-Joseph disease, and Parkin-PD (Rodríguez-Navarro et al. 2010; Sarkar et al. 2007; Seki et al. 2010; Zaltzman et al. 2020). However, the pharmacokinetics properties of trehalose are poor. The small intestine, kidney, and CNS expression of trehalose catalytic enzyme, trehalase, limits its bio-availability (Halbe e Rami 2019). Also, dietary trehalose results in digestive issues (Montalto et al. 2013) and may enhance the virulence of common nosocomial pathogen (Collins et al. 2018). These issues severely reduce the possibility of introducing trehalose into the clinical management of PD patients. In conclusion, boosting autophagy may be a powerful therapeutic strategy in LRRK2-PD, but a clinically relevant drug is still to be discovered.

## Experimental procedures

### Animals

All animal protocols were approved by the University of Trento and National Ministery of Health (IACUC 793/2016-PR). The murine *G2019S-LRRK2* BAC mice were previously described (Li et al., 2010) and have been backcrossed onto the C57BL/6J mice for >10 generations.

Animals were kept in a normal light/dark cycle (12 hours light/ 12 hours dark) and had free access to food and water. All procedures involving animals were approved by Institutional agencies (OPBA-Università degli Studi di Trento) and Italian Ministry of Health (Università degli Studi di Trento prot. n. 793/2016-PR).At 5 or 10 months, Wild-type and *G2019S-LRRK2* BAC male littermates were divided into trehalose and control groups. In the trehalose group, mice were offered 1% trehalose drinking water solution. The treatment solution was changed every week. In the control group, mice were given drinking water. Mice body weight was measured every week since the start of treatment. Only male mice were used in our experiments due to a strong gender effect (Pischedda et al., manuscript in preparation).

### Assays in post-mortem specimen

In total, 5 idiopathic, 4 G2019S and 5 control human brain cases were used in the study which were obtained from the archives of QSBB following appropriate local ethics committee approval. Demographic details of the cases have been described previously (Mamais et al. 2013). For biochemical analysis, 5μg of TBS soluble fraction and 10μg from TBS-5% SDS fractions of basal ganglia homogenates were run on 10% Bis-tris gels with MOPS as running buffer. Details of homogenisation and fractionation procedure is mentioned in a previous publication (Mamais et al 2013). Protein from gels were transferred onto PVDF membranes and probed with NSF (SYSY, mouse monoclonal) and beta actin (Sigma, mouse monoclonal) antibodies overnight with shaking. Following incubation with appropriate HRP conjugated secondary antibodies, blots were visualized by enhanced chemiluminescence (Pierce; UK) detection. Protein expression levels were determined by densitometry and NSF results normalised to beta-actin levels. For immunohistochemical evaluation, 8μm substantia nigra sections from control, 4 iPD and 2 G2019S-LRRK2 PD were dewaxed in xylene, and pre-treated with 95% formic acid for 10min to expose antigenic sites. Standard immunohistochemistry protocol was followed using NSF (1:200 mouse monoclonal; SY-SY) and alpha-synuclein (1: 300; rabbit polyclonal, Abcam) primary antibodies with o/n incubation at 4C. Double immunofluorescence was performed using tetramethyl rhodamine labelled secondary antibody for alpha-synuclein and NSF with the fluorescein signal amplification kit (Perkin Elmer, UK). Following adequate washes to remove background fluorescence, sections were mounted with Aquamount (Merck, UK). Control sections where primary antibody was omitted displayed no significant background staining. Fluorescent signals from sections were scanned using Leica fluorescent microscope (Leica CTR6000). Total alpha-synuclein positive LBs were counted from each section and the proportion of NSF+ve LBs were derived. Human brains were donated to the Queen Square Brain bank for neurological disorders, UCL Queen Square Institute of Neurology and stored under a research license No:12198.

### Behavioural tests

#### Spontaneous Motor Activity

Spontaneous motor activity was evaluated in an automated activity cage (43 cm × 43 cm × 32 cm) (Ugo Basile,Varese,Italy) placed in a sound-attenuating room as previously described (Ferri et al. 2004).Cumulative horizontal and vertical beam breaks were counted for 3 hours.

#### Balance Beam walking

The beam apparatus consists of 1 meter beams with a flat surface of 12 mm or 6 mm width resting 50 cm above the table top on two poles according to (Luong et al. 2011). A black box is placed at the end of the beam as the finish point. Nesting material from home cage is placed in the black box to attract the mouse to the finish point. A lamp (with 60 watt light bulb) is used to shine light above the start point and serves as an aversive stimulus. A video camera is set on a tripod to record the performance. On training days, each mouse crosses the 12 mm beam 3 times and then the 6 mm beam 3 times. The time required to cross to the escape box at the other end (80 cm away) is measured with a stopwatch. The stopwatch is started by the nose of the mouse entering the center 80 cm, and stopped when the animal reaches the end of the 80 cm. Once the mice are in the safe box, they are allowed some time (∼15 secs) to rest there before the next trial. The mice rest for 10 min in their home cages between training sessions on the two beams. On the test day, the time to cross each beam is recorded. The beams and box are cleaned of mouse droppings and wiped with towels soaked with 70% ethanol and then water before the next beam is placed on the apparatus.

#### Pole test

In the vertical pole task, the mouse was placed on a vertical wire-mesh pole with its head facing upwards. Mice were habituated to the task in 2 trials per day for 2 days. On test day (third day) mice were subjected to 5 trials: the total time taken to turn the body and to descend was recorded according to (Hickey et al. 2008). A cut-off of 60 sec was given. Data were shown as mean of 5 trials evaluated during the test day.

#### Rotarod

The rotarod apparatus (Ugo Basile, Biological Research Apparatus, Varese, Italy) was used to measure fore and hindlimb motor coordination and balance (Dauge et al. 2001). During the training period, each mouse was placed on the rotarod at a constant speed (12 or 32 rpm) for a maximum of 120 sec, and the latency (sec) to fall off the rotarod, within this time period, was recorded. Mice received four tests/trial each day for 4 consecutive days. The fourth test of each trial was evaluated for statistical analysis. The fifth day mice were submitted to Rotarod resistance at a constant speed (12 rpm) for a maximum of 30 min and the latency (sec) to fall was measured.

#### Novel object recognition

The novel-object recognition test was performed over a 3-day period in an open plastic arena (60 cm x 50 cm x 30 cm), as previously described (Pan et al. 2008). Animals were habituated to the test arena for 10 min on the first day. After 1 day habituation, mice were subjected to familiarization (T1) and novel-object recognition (T2). During the initial familiarization stage, two identical objects were placed in the centre of the arena equidistant from the walls and from each other. Each mouse was placed in the centre of the arena between the two objects for 20 min or until it had completed 30 s of cumulative object exploration. Object recognition was scored when the animal was within 0.5 cm of an object with its nose toward the object. Exploration was not scored if a mouse reared above the object with its nose in the air or climbed on an object. Mice were returned to the home cage after familiarization and then tested again after different delays (from 5 min to 24 h later). A novel object (never seen before) took the place of one of the two familiars. Scoring of object recognition was performed in the same manner as during the familiarization phase. From mouse to mouse the role (familiar or new object) as well as the relative position of the two objects were counterbalanced and randomly permuted. The objects for mice to discriminate consisted of white plastic cylinders and coloured plastic Lego stacks of different shapes. The arena was cleaned with 70% ethanol after each trial. An experimenter blind to the genotype group manually recorded the exploration times to the objects for each animal. Total time spent exploring the two objects during T1 and T2 was also calculated.

### Cell cultures

Neuron cultures were prepared from mouse cortexes obtained from embryonic day 15.5–16.5 wild-type or hG2019S mice (C57BL/6J) as previously reported (Pischedda e Piccoli 2015; Pischedda et al. 2018). Briefly, after brain dissection the cortexes were mechanically dissociated after 15 min incubation with 2,5% Trypsin (Euroclone) at 37 °C in agitation. The resulting cells were counted and cryopreserved or directly plated on previously poly-D-lysine (Sigma) coated wells or cover slips, according to the desired density. High density (750–1000 cells/mm2) neuron cultures were plated on 12-well plastic tissue culture plates (Iwaki; Bibby Sterilin Staffordshire, UK) and medium-density (150–200 cells/mm2) onto 12mm diameter cover slips inserted into 24-well plastic tissue culture plates (Iwaki). The cells were grown in Neurobasal at 37°C and 5% CO_2_. Neuronal transfection was carried out using Lipofectamine 2000 (Life Technologies) following the manufacture’s instruction. The cells were treated with LRRK2 kinase inhibitor, GSK-2578215A (Tocris Bioscience, Bristol, UK), MG132 (Tocris Bioscience, Bristol, UK) and Trehalose (Sigma) by addition to culture media at the concentrations indicated through the text. Human embryonic kidney cells (HEK293T) were cultured in DMEM complete: Dulbecco’s modified Eagle’s medium (DMEM, Euroclone) supplemented with 10% fetal bovine serum (FBS, Euroclone) at 37°C and 5% CO2. HEK293T were transiently transfected using linear polyethylenimine (PEI, Polysciences) with ratio DNA: PEI 0.8:100. 4 µg of DNA were dissolved in 0.5 ml of PEI solution and vortexed for 20 sec. The DNA-PEI mix was incubated for 10 minutes at room temperature (RT) and added directly to the cells. The medium was changed after 24h and the cells were lysed 48h post transfection. Lentiviruses were produced by transient transfection of HEK293T cells according to standard protocols (Wiznerowicz e Trono 2003). Primary cortical cultures were transduced at DIV4 with lentiviruses at multiplicity of infection 3 (MOI3). The 3-(4,5-dimethylthiazol-2-yl)-2,5-diphenyltetrazolium bromide (MTT) assay was performed to measure culture viability as described (Mosmann 1983). High density (750–1000 cells/mm2) neuronal culture were infected at DIV 4 and processed for MTT assay at DIV16. Briefly, MTT ((3-(4, 5-dimethylthiazolyl-2)-2, 5-diphenyltetrazolium bromide-Sigma Aldrich) stock solution 20X (5mg/ml in MilliQ water) was diluted in neuronal complete media and cells were were incubated with MTT mix for 30 minutes at 37°C with 5% CO2. After incubation, media was discarded, cells were resuspended in DMSO to dissolve the intracellular purple precipitates and the absorbance was measured at 570 nm with the Plate reader Infinite 200 PRO (Tecan Life Sciences). Toxicity was calculated as 1-cell viability, being cell viability expressed as fold over control.

### iDA culture

Wild-type, G2019S and G2019S corrected Human iPSCs derived Neuronal Precursors Cells (NPCs) were obtained from Dr. Deleidi lab and cultured as previously reported (Reinhardt et al. 2013). Briefly, the cells were expanded in 50:50 DMEM F12 and Neurobasal medium with: 1% P/S, 1% B27 (without Ascorbic acid), 0,5% N2, 1%L-glutamax, 150 µM Ascorbic acid, 3 µM CHIR, 0.5 µM Purmorphamine (PMA) (NPC expansion medium). Media was changed every other day and cells were passed every week after 10 min incubation with Accutase (200ul/12 well). The cells were re-plated at a 1:10 ratio in 12-well plates pre-coated with Matrigel (1:30 from Stock) in ‘NPC expansion medium’ supplemented with 10 μM ROCK inhibitor for the first day after splitting. After the second passage, when the cells reached 80% of confluency, the media was changed to Ventral CNS differentiation medium (50:50 DMEM F12 and Neurobasal medium with: 1% P/S, 1% B27 (without Ascorbic acid), 0,5% N2, 1%L-glutamax, 100 ng/mL FGF8, 200 µM of Ascorbic acid and 1µM PMA. When confluent the cells were split with Acutase and re-plated in Matrigel coated 6 wells plate at a 1:1 ratio. The media was changed every other day. After 10 days the Ventral CNS differentiation medium I was changed to maturation medium (50:50 DMEM F12 and Neurobasal medium with: 1% P/S, 1% B27 (without Ascorbic acid), 0,5% N2, 1%L-glutamax, 10 ng/mL BDNF, 10 ng/mL GDNF, 1ng/mL TGF-b3, 200 µM ascorbic acid, 500 µM dbcAMP and 0,5 µM PMA). After 48h the media was changed to dopaminergic differentiation medium (50:50 DMEM F12 and Neurobasal medium with: 1% P/S, 1% B27 (without Ascorbic acid), 0,5% N2, 1%L-glutamax, 10 ng/mL BDNF, 10 ng/mL GDNF, 1ng/mL TGF-b3, 200 µM ascorbic acid, 500 µM dbcAMP). The cells were kept in dopaminergic differentiation medium until the end of the experiment and the medium was changed every 2-3 days.

### Immunohistochemistry

Six and 12 months old mice were anaesthetized and transcardially perfused with PBS 1X followed by 4% paraformaldehyde in PBS 1X. Brains were dissected, postfixed for 2h by immersion in the same fixative and then cryoprotected by immersion in 30% sucrose solution in PBS 1X for 24 h at 4°C. Brains were included in OCT and stored at −80°C until processing. 14 µm brain coronal sections were obtained after serial sectioning on Leica cryostat and mounted on Polysine Slides (Thermo Fischer Scientific). The slices were saturated in 2,5% BSA, 10% NGS, 0,2% Triton X-100, PBS 1X, for 1 hour at RT and incubated with the primary antibody O/N at 4°C. After 3 washes in PBS 1X – 0,2% Triton X-100 the sections were quenched for 15 min in PBS 1X plus 0,3% H_2_O_2_at RT, then washed with PBS 1X. The sections were incubated for 30 min with the secondary biotinylated antibody at RT. The slices were then rinsed in PBS1X and incubated for 30 minutes with Vectastain ABC kit 1:1 mix (Vectorlab), washed 2 times in PBS 1X – 0.2% Triton X-100, washed once in PBS1X, and developed with DAB quanto mix (Thermo Scientific) for 30 sec. The stained sections were rinsed in PBS 1X stained in Hematoxylin solution, Gill No.2 (Sigma Aldrich) for 2,5 minutes and then washed for 2 minutes with tap water. Sections were incubated with differentiation solution (70% ETOH plus 0,3% HCl) for 10 seconds, washed with tap water and then incubated for 30 seconds with Scott’s Tap water Substitute (Sigma Aldrich) for blueing. Sections were then dehydrated, mounted using DPC mounting reagent (Sigma) and kept at RT until acquisition with a Zeiss Axio Imager M2 equipped with 40X objective.

### Immunofluorescence

For the immunostaining experiments, neurons were fixed in 4% paraformaldehyde and 4% sucrose at room temperature or 100% methanol at −20°C. mCherry positive neurons were randomly chosen for quantification in at least four independent experiments for each condition. Cover slips were mounted with prolonged reagent (Life Technologies) and observed with Zeiss Observer Z1 microscope equipped with an Apotome module. The obtained images provide an axial resolution comparable to confocal microscopy(Garini, Vermolen, e Young 2005; Schaefer, Schuster, e Schaffer 2004). Images were acquired with AxioObserv Z1 microscope equipped with Apotome module using a plan-Apochromat 63x/1.40 Oil objective, pixel size 0,102 mm x 0.102 mm. Acquired images were analyzed with ImageJ software using NeuronJ plugin (Meijering et al. 2004).

### Polyclonal phospho-specific NSF antibody production and purification

Affinity-purified anti-P-Thr645-NSF polyclonal antibody was made in a NZW female rabbit (Envigo) by immunization against a single peptide (amino acids 631–656: KKAPPQGRKLLIIGpTTSRKDVLQEME), encompassing the LRRK2 phosphorylation site of NSF, conjugated via an N-terminal Cys residue to maleimide-activated KLH (keyhole limpet haemocyanin, Sigma-Aldrich). Rabbit was injected subcutaneously four times at 3-week intervals, with 0.5□mg of phosphopeptide-protein conjugates in PBS emulsified with Freund’s adjuvant (1:1 v/v). After the last immunization, blood was collected and antisera was purified using an immobilized peptide affinity resin (Sulfo Link Coupling Gel, Thermo) according to manufacturer’s instructions. Briefly, a shorter peptide (LIIGpTTSRKD) was conjugated via an N-terminal Cys residue and coupled to the resin. Fractions were eluted with Glycine Buffer (0.2 M, pH 2.5) into Tris buffer (1 M, pH 8.5) solution for pH neutralization. Protein eluted fractions (as determined by Abs at 280 nm) were pooled and concentrated by ultrafiltration (Amicon Ultra-15 30KD, Millipore). All synthetic peptides were synthesized, purified by RP-HPLC, and confirmed by mass spectroscopy at CRIBI (University of Padova) as previously described (Ottaviani et al. 2017). All animal protocols were approved by the University of Padova and National Ministery of Health (IACUC 906/2016-PR).

### Biochemical assays

#### Proteinase K digestion

Proteinase K digestion of mouse brain lysates was performed on samples prepared from 6 months mouse LRRK2 wild-type and LRRK2 G2019S brains. Brains were solubilized by mechanical homogenization of 1g of tissue in lysis buffer (150 mM NaCl, 2 mM EDTA, 50mM Tris-HCl, 1% NP-40 and 0.25% sodium deoxycholate, pH 7.4) using a pestle and incubated 1h at 4°C. Next brain lysates were centrifuged at 14,000xg for 10 min at 4°C to collect supernatant and samples were quantified following Bradford’s method. Proteinase K was added to 50 µL of sample (previously prepared at 1µg/µL concentration) at different final concentration ranging from 0.3-0.5 µg/mL and incubated at 37°C for 25 minutes under shaking. Digestion reactions were stopped by addition of the inhibitor PMSF (4mM) and samples (undigested and exposed to Proteinase K) were prepared in sample buffer 5X for Western-blotting analysis.

#### Proteasome activity

Proteasome activity in mice brain tissue was analysed following the 20S Proteasome activity assay kit protocol by Millipore (APT280). The assay detects the fluorophore 7-Amino-4-methylcoumarin (AMC) after cleavage from the labelled substrate LLVY-AMC. Briefly, 3 µg of protein from each sample was incubated for 1 h at 37°C with proteasome labeled substrate in 96 dark multiwells. The sensibility of the assay was tested performing serial dilution of proteasome positive control (1:4 to 1:256) of the stock solution in 1X Assay buffer. Fluorescence data was collected using a PE Biosystems CytoFluor 400 plate reader using a 380 nm excitation and 460nm emission filters.

#### Filter retardation assay

Filter retardation assay were performed as described in (Perez Carrion et al. 2018). Briefly, cell cultures were rinsed once in cold PBS 1X and mechanically detached in PBS 1 X plus protease and phosphatase inhibitor cocktails Calbiochem (40 µl/well 12 wells plate). Cell samples were sonicated 3 times with 1s pulse and 1s stop sequence at 10% pulse amplitude. Brain extracts from 6 months mouse LRRK2 wild-type and LRRK2 G2019S variant were prepared by mechanical homogenization of 1 gr of tissue in solubilization buffer (150 mM NaCl, 2 mM EDTA, 50mM Tris-HCl, pH 7.4) using a pestle and incubated 1h at 4°C. Then brain lysates were centrifuged at 14,000xg for 10 min at 4°C to collect supernatant. Protein amounts were quantified following Bradford’s method. Ten µg of each sample were diluted to 120 µl in PBS 1X. The samples were immobilized on methanol equilibrated cellulose acetate membrane through a dot blot void system (BioRad).

#### Streptavidin pull down

Streptavidin pull down was performed as described in (Pischedda et al. 2014). Briefly, HEK293T cells were transiently transfected and treated as indicated trough the text. The cells were then resuspended in 5ml ice cold PBS 1X, centrifuged at 150xg for 5 min and lysed in 1 ml of 50mM HEPES pH 7.4, 150mM NaCl, 0.5mM EDTA, 0.5% Triton X-100, protease inhibitors (Calbiochem) and phosphatase inhibitor cocktails II and IV (Calbiochem), after incubation for 1 h at 4°C in rotation. The cell lysate was recovered after 10 min centrifugation at 15000xg at 4°C and 10% of supernatant kept as input. The remaining supernatant was incubated with 25 µl of Streptavidin resin (IBA) for 1 hour at 4°C in rotation. After three washes in 0.5 ml of 50mM HEPES pH 7.4, 150mM NaCl, 0,2% Triton X-100, the bound protein was eluted in 60 µl of Laemmli buffer 1X after 10 min incubation at 52°C.

#### Solubility assay

HEK293T cells were cultivated in 12 well plates and transiently transfected and treated as indicated trough the text. Cells were rinsed with cold PBS1X and then lysed in Triton-X100 buffer at 4°C (50mM HEPES pH 7.4, 150mM NaCl, 1 % Triton X-100, protease inhibitors (Calbiochem) and phosphatase inhibitor cocktails II and IV (Calbiochem)). The cell lysate was kept on ice and briefly vortexed every 10 minutes for 3 times, then centrifuged at 16.000g for 10 minutes at 4°C. The Supernatant was collected as the detergent-soluble fraction and the pellet constituted the insoluble fraction. The pellet was resuspended in 75 ul of Laemmli sample buffer 2X, whereas the same volume of Laemmli sample buffer 5X was added to the soluble fraction.

#### Protein purification

HEK-293T cells were lysed in lysis buffer [20 mM Tris–HCl, pH 7.5, 150 mM NaCl, 1 mM EDTA, 2.5 mM Na4P2O7, 1 mM beta-glycerophosphate, 1mM NaVO4, Protease Inhibitor Mixture (Sigma-Aldrich)] by using a needle. Cleared lysates (1 ml) were incubated with 30 µl of anti-FLAG-M2-agarose beads by rotating overnight at 4°C. Resin complexes were washed with different buffers WB1 (20mM Tris–HCl pH7.5, 500mM NaCl) twice, WB2 (20mM Tris–HCl pH7.5, 350mM NaCl) twice, WB3 (20mM Tris–HCl pH7.5, 150mM NaCl) six times. Proteins were eluted in WB3 with 150 ng/µl of 3× FLAG peptide) for 30 min at 4°C with shaking. Purified proteins were resolved by SDS–PAGE and stained with Coomassie Brilliant Blue to verify protein purity. Protein content was calculated by densitometry, using a standard curve with bovine serum albumin (BSA).

#### ATPase assay

NSF ATPase activity was quantified using the Malachite Green Assay by measuring the release of inorganic phosphate (Pi) due to the ATP hydrolysis with spectrophotometer as previously described (Belluzzi et al. 2016). 216 nM NSF was incubated with different ATP concentrations and 2 mM MgCl2 for 2h a 37°C. The samples were mixed with Malachite Green stock solution and the absorbance measured at 640nm. The values of absorbance were then converted into μmol of free Pi in solution using a standard curve. Kinetic constants were obtained by data fitting with the Michaelis-Menten kinetic model Y□=□Vmax*S/(Km□+□[S]).

#### Size Exclusion Chromatography (SEC) and dot blot analysis

SEC was performed as previously described (Belluzzi et al. 2016). Briefly, Strep-FLAG NSF WT and T465D transfected HEK-293T cells were lysed in 0.5ml of lysis buffer containing 0.06% (v/v) Triton X-100 and centrifuged. Cell lysates were separated on a Superose 6 10/300 column (Ge Healthcare, Waukesha, WI, USA) pre-equilibrated with 20mM Tris–HCl pH7.5, 150mM NaCl and 0.06% (v/v) Triton X-100. The flow rate used was 0.5ml/min. Fractions of 0.25ml were collected and spotted onto a nitrocellulose membrane and analyzed by dot blot. Densitometric analysis was performed by using ImageJ program.

#### Western-blotting

Protein samples were resolved on to 10-15% Tris-glycine polyacrylamide gels in SDS/Tris-glycine running buffer. Solubilized proteins were transferred to nitrocellulose or polyvinylidenedifluoride (PVDF) membranes in blotting buffer containing 10% methanol. The membrane was saturated for 1 h at RT in 5% nonfat dry milk in Tris-buffered saline and 0.1% Triton (TBS-T) and then incubated with the primary antibody in saturating solution for 90 min at RT. The unbound primary antibody was sheets was washed out in TBS-T (3×10 min) at RT and the membrane was incubated for 1 h at RT with horseradish peroxidase-conjugated anti-mouse or anti-rabbit IgGs. After 3 washes in TBS-T 10 min/each at RT, the immunoreactive proteins were visualized using enhanced chemioluminescence plus (ECL+, GE Healthcare, Little Chalfont, England). Densitometric analysis was carried out using Image J software.

#### Structure analysis

The Cryo-EM structure of NSF hexamer was downloaded from the PDB database, and the structural analysis was performed using the program PyMOL (*http://www.pymol.org/*). All the structural diagrams were prepared using the program PyMOL.

### Statistical analysis and guidelines

All data are expressed as mean ± standard error of the mean (S.E.). The normality of data distributions was determined using the D’Agostino and Pearson omnibus normality test, followed by an unpaired Student’s t test, ANOVA followed by Tuckey’s post-hoc test or two-way ANOVA followed by Bonferroni or Student’s t post-hoc test as appropriate. The indication of number of experiment (n) and level of significance (p) are indicated throughout the text. All methods were performed in accordance with the relevant guidelines and national regulations.

### List of expressing vectors

HA-Ubiquitin (pcDNA3.1), HA-K48R Ubiquitin (pcDNA3.1); HA-K63R Ubiquitin (pcDNA3.1) HA-K29,48,63R Ubiquitin (pcDNA3.1) (kind gifts by Dr. Simona Polo, IFOM);

LRRK2 K1906M GFP (pcDNA3.1), LRRK2 G2019 GFP (pcDNA3.1) [described in (Gloeckner et al. 2006);

Strep-FLAG hNSF WT (pcDNA3.1), Strep-FLAG hNSF T645A (pcDNA3.1) [described in (Belluzzi et al. 2016)], Strep-FLAG hNSF T645DE (pcDNA3.1) and Strep-FLAG hNSF T645D (pcDNA3.1) generated by site directed mutagenesis;

HA hNSF (pCMV HA-2) (kind gift by prof. Dario Alessi);

Strep-FLAG hNSF WT (pULTRA HOT) and Strep-FLAG hNSF T645D (pULTRA HOT) generated via cloning into NheI-BclI sites

### List of primary antibodies

NSF (123 002-Synaptic System); alpha-Synuclein (D37A6-Cell Signaling); LC3 (NB100-2331-Novus Biologicals); p62 (ab109012-ABCAM); b-ACTIN (sc-47778 – Santa Cruz Biotechnology); FLAG (sc-166355-Santa Cruz Biotechnology); HA (Cat. #05-904-Merck Millipore); cleaved-caspase-3 (9H19L2-Invitrogen); TH (AB152-Merck Millipore); b3-Tubulin (ab18207-ABCAM); Synapsin-1 (D12G5-Cell Signaling); RAB 3a (107 111-Synaptic System); LRRK2 phospho S935 (UDD2 10(12) ab133450 – ABCAM); LRRK2 (MJFF2 (c41-2) ab133474 – ABCAM); LRRK2 phospho S1292 (MJFR-19-7-8 ab203181 – ABCAM)

### List of secondary antibodies

Anti Rabbit IgG (H+L), made in goat, Biotinylated (BA-1000-Vector Laboratories); Anti Mouse IgG (H+L), made in horse, Biotinylated (BA-2000-Vector Laboratories); Alexa Fluor® 488 AffiniPure Goat Anti-Rabbit IgG (H+L) (111-545-144 - Jackson ImmunoResearch); Goat IgG anti-Rabbit IgG (H+L)-TRITC (111-025-003 - Jackson ImmunoResearch); Peroxidase AffiniPure Goat Anti-Mouse IgG (H+L) (115-035-146 - Jackson ImmunoResearch); Peroxidase AffiniPure Goat Anti-Rabbit IgG (H+L) (115-035-003 - Jackson ImmunoResearch)

## Author Contributions

F.P., M.D.C, L.P., M.S., A.B., and M.P.C. performed experiments. O.M., M.M., Li.P., E.G., R.B., M.E.S., and G.P. analysed data and wrote the paper.

## Acknowledgments

G.P. is supported by Fondazione Telethon (grant TDPG00514TA), MIUR (PRIN-2017ENN4FY), and Fondazione Cariplo (project 2019-3415). F.P. received support by Fondazione Caritro (project 2019.0230). This work was supported by Fondazione Cariplo (grant 2011-0540) to G.P. and E.G. and Fondazione Telethon (grant GGP12237) to G.P., E.G. and M.M. G.P. is grateful to the Michael J. Fox Foundation, the FIRB program (grant RBFR08F82X_002) and Fondazione Grigioni per il morbo di Parkinson. R.B. is funded by the Reta Lila Weston Trust and the British Neuropathological Society. We gratefully thank Prof. Matthew Farrer and Dr. Heather Melrose for providing LRRK2 hG2019S BAC mice, to Marzia Indrigo for excellent technical advice and Giuseppe La Tona for support in animal handling.

---

**Supplementary figure 1. hG2019S mice present proteinaceous aggregates**. Wild-type and hG2019S mice were processed for imaging analysis at 6-month. Brain sections were stained with anti-LC-3 antibodies and counterstained with hematoxylin to visualize nuclei. Scale bars = 50μm. We reported a peculiar LC3-immunoreactivity in the nigra, cortex, and hippocampus specimen prepared from hG2019S mice brain. Insets highlight differences in LC3 immunoreactivity in neurons (2X higher magnification). We noticed intense LC3 staining surrounding *bona fide* pale bodies (indicated by the asterisk) in nigra (e-f), cortex (q-r), and hippocampus (w-x).

**Supplementary figure 2**. Wild-type and hG2019S mice were processed for imaging analysis at 6-month. Brain sections were stained with anti synuclein (A) or anti Rab3A (B) antibodies and counterstained with hematoxylin to visualize nuclei. Scale bars= 50μm.

**Supplementary figure 3**. We performed a biochemical analysis of cortical (Cx), hippocampal (Hi), and striatal (St) samples harvested from 6 months old wild-type and hG2019S mice brains (A). The graphs report LRRK2 phosphorylation at Ser935 fold-over total LRRK2 (B), relative LRRK2 (C), and NSF (D) amount. Data are shown as mean ±SEM, n=8. We performed a western-blot analysis to profile the maturation of G2019S or gene-corrected neural precursor cell to dopaminergic neurons. Anti-nestin antibody stains immature proliferating cells while TH mature DA-neurons. I: expansion media; II: ventral CNS neuron differentiation medium; III maturation medium; IV: differentiation medium (E). Immunofluorescence characterization of terminally differentiated NPC shows the expression of β-III-tubulin, synapsin I, and TH proteins. Scale bars = 10μm (F).

**Supplementary figure 4**. Validation of a polyclonal anti-P-Thr645 NSF antibody. HEK293 cells were transfected with LRRK2-G2019S and FLAG-tagged wild-type NSF protein. 48h post-transfection cells were treated with increasing doses of MLi-2 for 4 hours, then lysed and analyzed (40µg of proteins) by immunoblotting (A). HEK293 cells were transfected with FLAG-tagged wild-type NSF protein or NSF T645A or the phosphomimetics NSF T645E and T645D variant together with GFP or LRRK2-G2019S. Upon solubilization, lysates (40µg of proteins) were subjected to immunoblotting (B). Affinity-purified anti-P-Thr645-NSF was used for immunoblot at 1:1000 dilution (in 3% BSA in TBS-T), and in the presence of an excess (5X) of non-phosphorylated peptide antigen (KKAPPQGRKLLIIGTTSRKDVLQEME). LRRK2 phosphorylation influences NSF aggregation. We assayed the solubility profile of NSF in samples prepared from N2A cell over-expressing LRRK2 K1906M or LRRK2 G2019S variants and treated with vehicle (DMSO) or the LRRK2 kinase inhibitor GSK2578215A (GSK, 0.2 mM, 18 hours) (C). The graphs report the amount of NSF present in the Triton-X100 insoluble fraction, expressed as fold-over total NSF. Data are shown as mean ±SEM, n=7. ** p<0.01 versus wild-type, same treatment; # p<0.05 versus G2019S DMSO (D). NSF aggregates in G2019S LRRK2 cellular models independently from proteasome activity. Wild-type and hG2019S DIV14 cortical neurons were treated with 100nM MG-132 or vehicle (DMSO) for 48 hours and then assayed by filter retardation assay to isolate high molecular weight (HMW) form of NSF or by dot-blot to measure total NSF protein. NSF appears in HMW aggregates in wild-type neurons upon proteasome impairment and in vehicle-treated hG2019S neurons (E). The graphs report NSF aggregation expressed as HMW fold-over total NSF. Data are shown as mean ±SEM, n=8. *** p<0.001 versus wild-type, same treatment, ## p<0.01 versus DMSO, same genotype (F). We analyzed 20S proteasome activity in brain samples prepared from 6- or 12-months old mice. The graph reports relative fluorescence at 520-530 nm and is expressed as mean ±SEM, n= 4 (G).

**Supplementary figure 5**. Inorganic phosphate (Pi) generated by ATP hydrolysis in the presence of NSF wild-type and T645D mutant was measured with the Malachite Green Assay at 120min. Data are represented as means ± S.E.; n=3, **=p<0.01 Student T-test (A). Kinetic constants were obtained by data fitting with the Michaelis-Menten kinetic model Y□=□Vmax*S/(Km□+□[S]). Vmax NSF wild-type=3.6 µmol/min NSF T645D=4 µmol/min; Km NSF wild-type=188 µM Km NSF T645D=180 µM. Extra sum-of-squares F test was used for statistical analysis; n=3 ***p<0.001 (B).

HEK293 cell over-expressing Strep-FLAG-NSF wild-type or phosphomimetic T645D isoform and treated with vehicle (DMSO) or MG-132 (10 μM, 18 hours) were analyzed by filter retardation assay to isolate high molecular weight (HMW) form of NSF or by dot-blot to measure total NSF protein (C). The graphs report NSF aggregation expressed as the ratio of HMW over total NSF optical density. Data are shown as mean ±SEM, n=5. ** p<0.01 versus wild-type, same treatment (D). Size exclusion chromatography fractions of HEK293T expressing ectopic Strep-FLAG-NSF wild-type or T645D alone spotted onto nitrocellulose membrane and probed with anti-flag antibody. Theoretical molecular weight are V0 at fraction 8.5, 669kDa at fraction 12, 449kDa at fraction 13 (E). The graph reports the intensity of each dot (fraction) normalized by the integrated intensities. The column void volume is 7.5 ml, n=3 (F). We transfected in HEK293T cells myc-ubiquitin wild-type, K48R, K63R, or K29/48/63 R together with Strep-FLAG-NSF WT or T645D. We purified on streptavidin-beads NSF protein and assessed ubiquitination level by western-blotting with anti-myc antibody. Arrowheads indicate putative NSF low molecular weight fragments detected by anti-FLAG antibody (G). The graphs H, J report NSF ubiquitination calculated as anti-myc optical density normalized versus total NSF wild-type (H) or T645D (J) yield and expressed as fold-over ubiquitin wild-type condition. The graphs I, K and NSF fragmentation, expressed as the ratio of low molecular weight NSF fragments over high molecular weight NSF wild-type (I) or T645D (K) optical density and expressed as fold-over ubiquitin wild-type condition. Data are shown as mean ±SEM, n=7. *, p<0.01, ** p<0.01 versus ubiquitin wild-type, one-sample T-test.

**supplementary figure 6**. Wild-type and hG2019S cortical neurons were infected at DIV4 with mCherry expressing viruses or viruses co-expressing mCherry and wild-type Strep-FLAG NSF (NSF WT) or Thr645Ala Strep-FLAG NSF (NSF T645A). Cells were chronically treated from DIV4 to DIV14 with DMSO or GSK2578215A (0.2 μM, every two days). Cells were then solubilized and processed for western-blotting to assess ectopic NSF expression, LRRK2 phosphorylation, and total LRRK2 level (A). DIV13 Wild-type and hG2019S cortical neurons were treated with vehicle (water) or trehalose (100 mM, 1 day) alone or in combination with NH4Cl (5mM, 2 hours) and then processed for western blotting to evaluate the induction of autophagy, indicated by the appearance of LC3II band (B). The graph reports the LC3II/LC3I ratio. Data are shown as mean ±SEM, n=8. *** p<0.001 versus water (C). We assayed by filter retardation assay the presence of NSF in high molecular weight aggregates in samples prepared from HEK293 cell over-expressing NSF wild-type and treated with vehicle (DMSO) or MG-132 (10 μM, 18 hours) alone or in combination with trehalose (100 mM, 18 hours) (HMW) as well as total NSF expression (total) via dot-blot on a nitrocellulose membrane (D). The graphs report NSF aggregation expressed as ratio of HMW over total NSF optical density. Data are shown as mean ±SEM, n=8. ** p<0.01 versus DMSO; # p<0.05 versus control treatment (E). We assessed by filter retardation assay NSF aggregation in brain specimens prepared from 6-month wild-type and hG2019S mice treated with trehalose (1% in drinking water, 1 month). Trehalose treatment reduces HMW NSF forms in cortical specimens obtained from 6 months mice (E). The graphs report NSF aggregation expressed as the ratio of HMW over total NSF optical density. Data are shown as mean ±SEM, n=6. * p<0.05 versus wild-type, same treatment, ## p<0.01 versus water, same genotype (G). Trehalose treatment does not influence mice weight. The graph reports mice weight along with the 4 week treatment. Data are shown as mean ±SEM, n=10 (H).

**supplementary figure 7**. Wild-type and hG2019S mice were treated starting at 10 months with trehalose (1% in drinking water) and profiled for motor and cognitive abilities at 12 months. In detail, we measured spontaneous motor activity in terms of the umber of horizontal (A) and vertical (B) counts in 3 hours, time to cross a 6-mm or 12-mm width beam (C), time to reach the ground from a supra-elevated platform (D), time spent on rotarod running at 12 rpm (E) or 32 rpm (F), over 4 trials (T_1_-T_4_), total resistance on 12 rpm running rotarod (G) and ability to recognize a novel object (H). Data are shown as mean ±SEM; n=7-18. * p<0.05, ** p<0.01, *** p<0.001,****p<0.0001 versus wild-type, same treatment, $ p<0.05, $$$ p<0.001 versus water, same genotype.

