## supplementary figures 1-7 for "LRRK2 G2019S kinase activity triggers neurotoxic NSF aggregation"

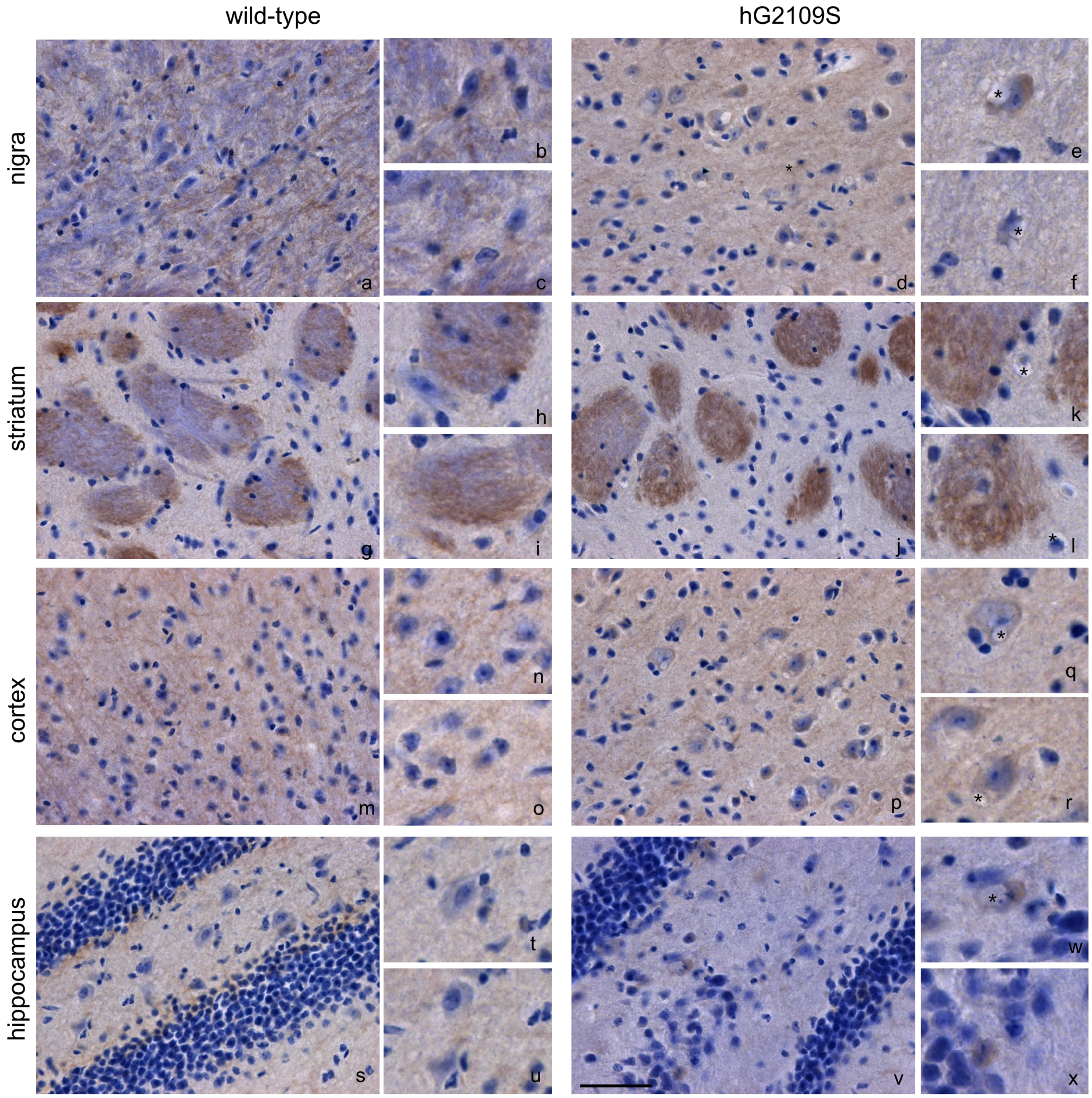


supplementary figure 1

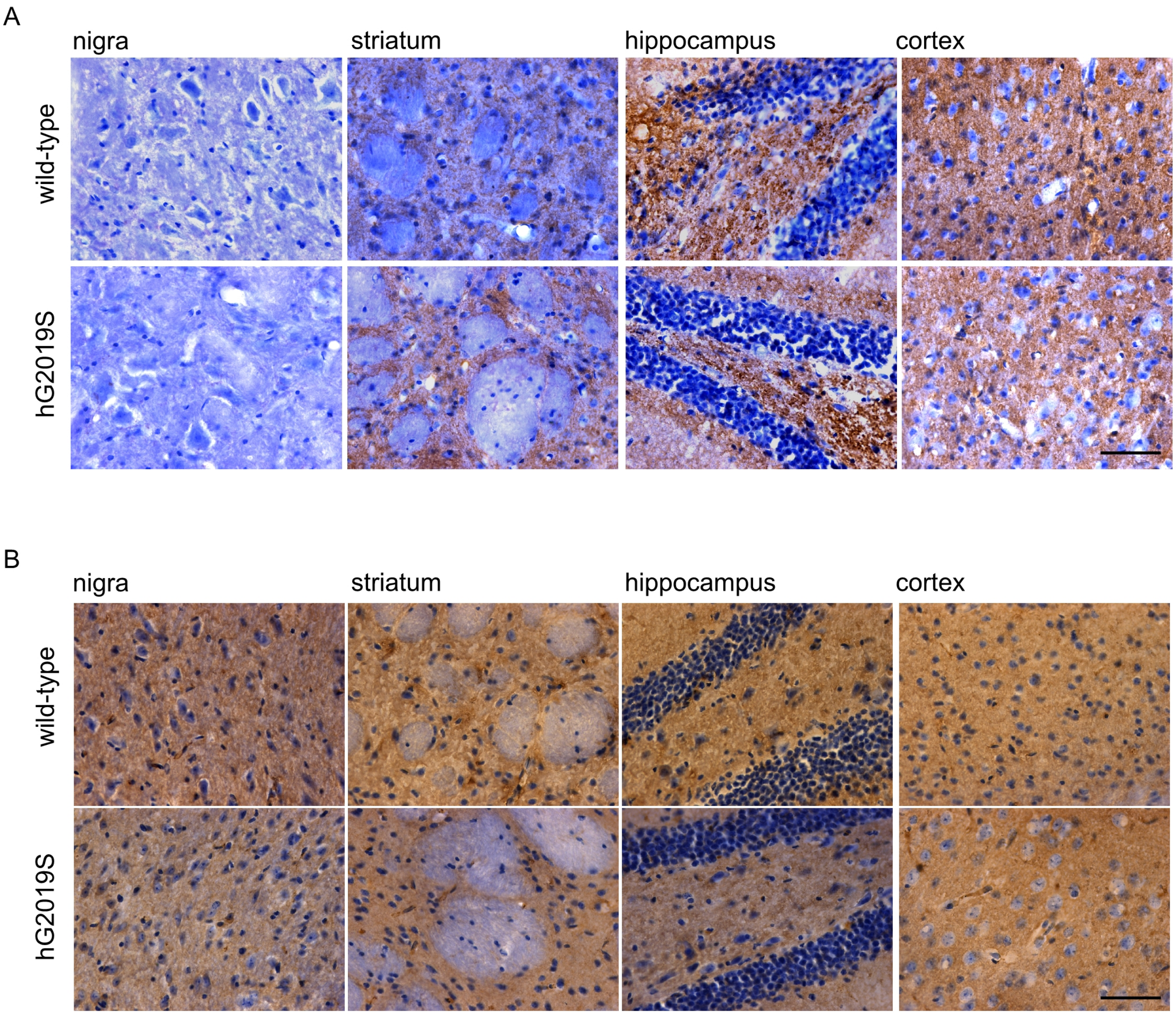


supplementary figure 2

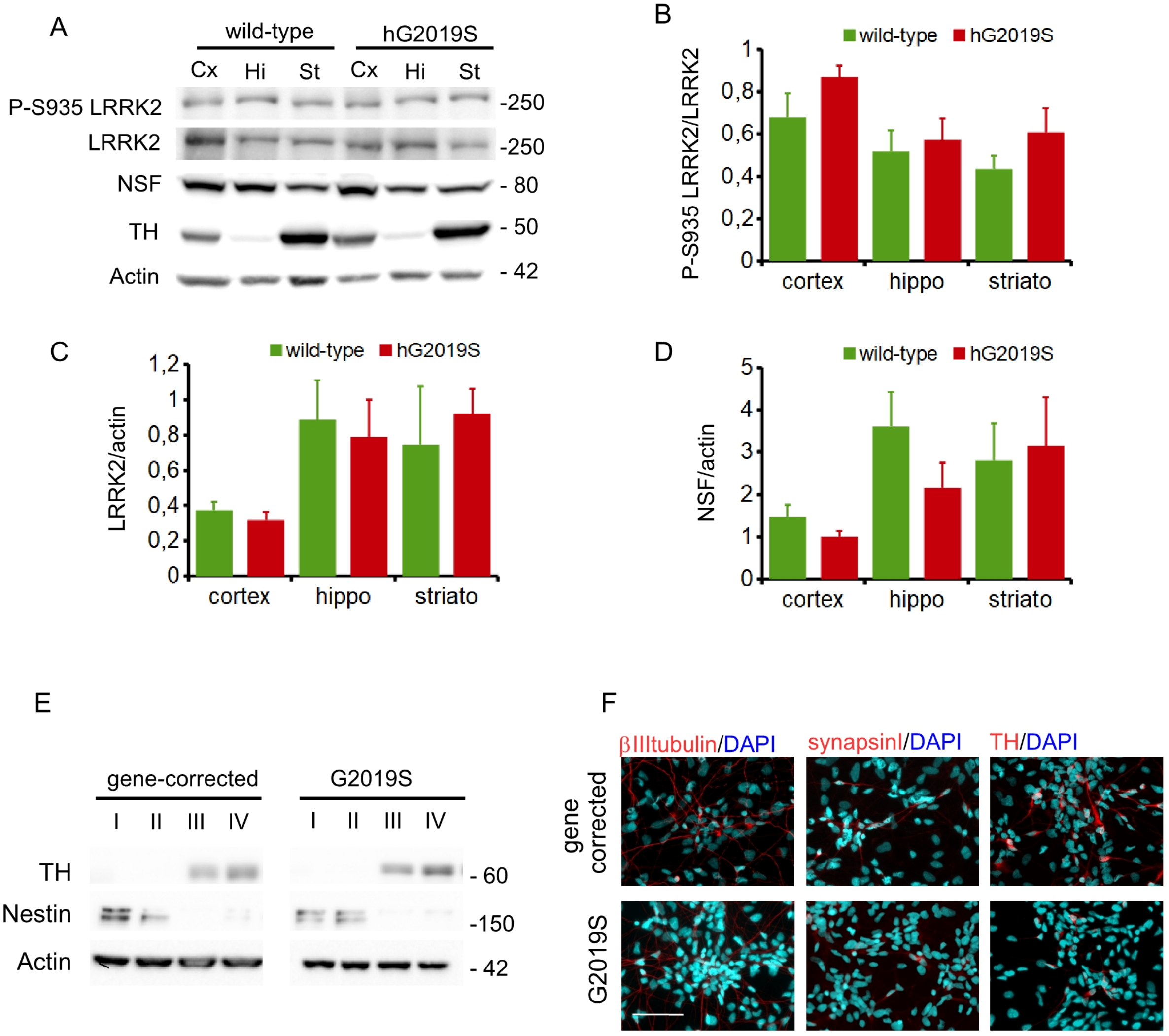


supplementary figure 3

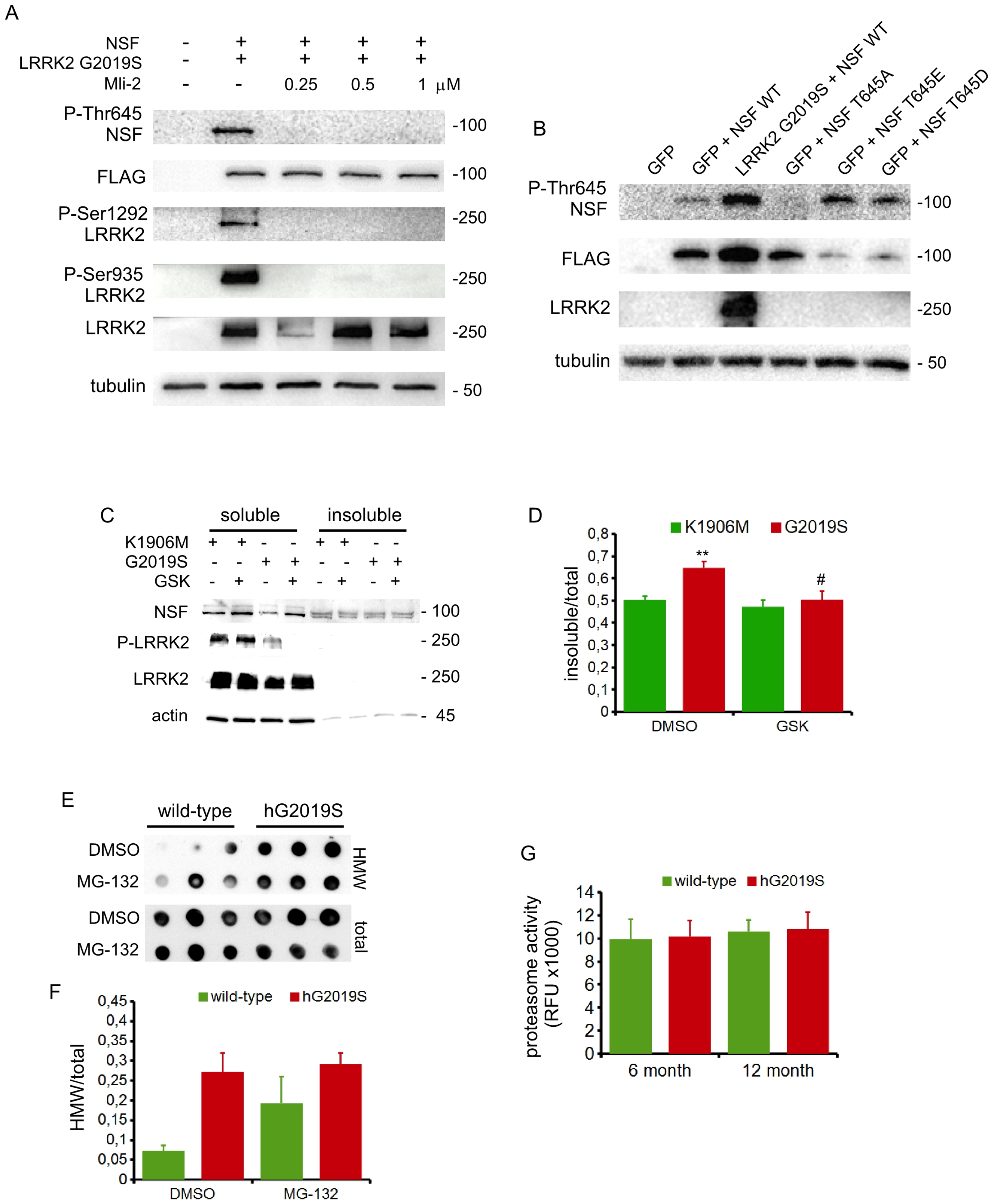


supplementary figure 4

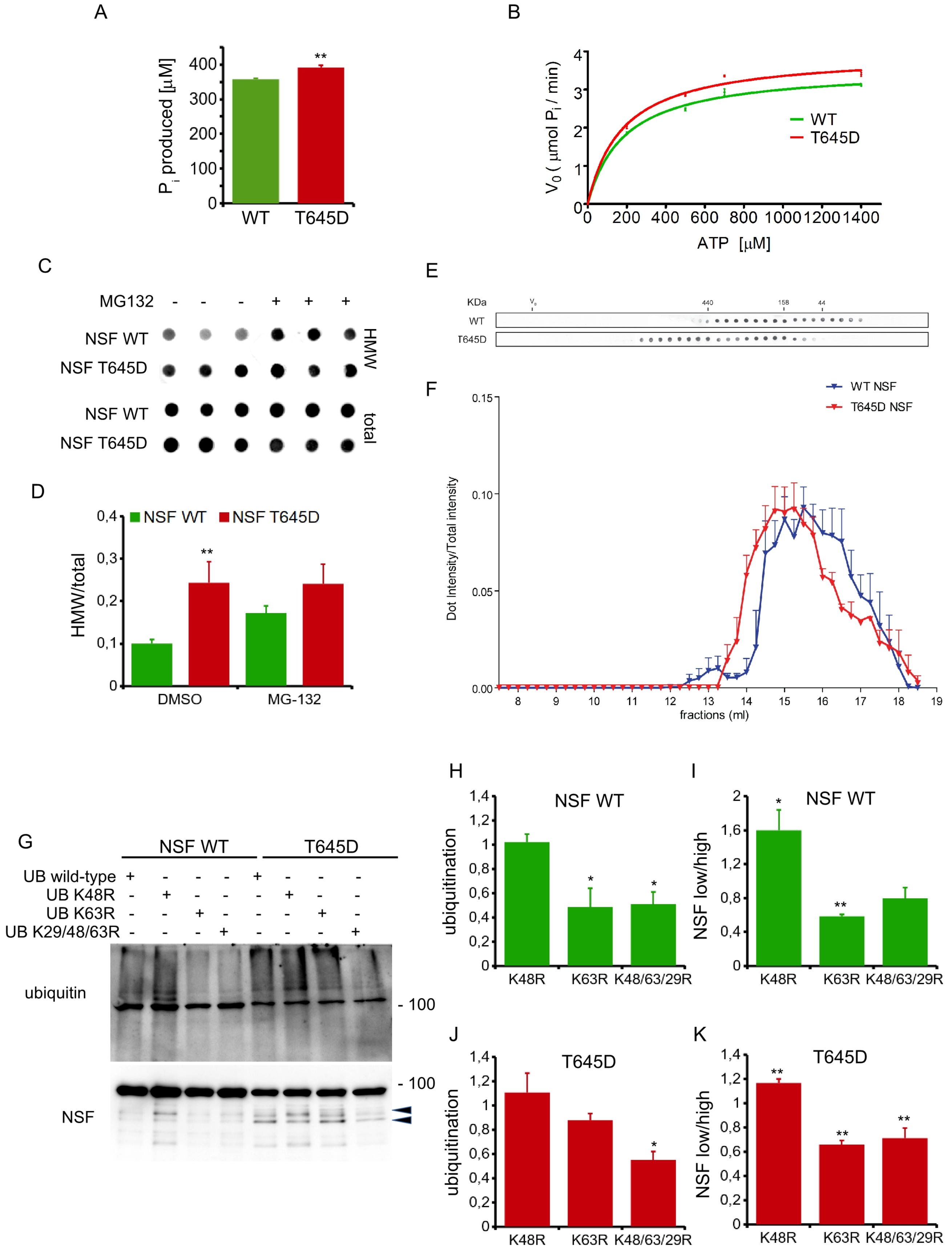


supplementary figure 5

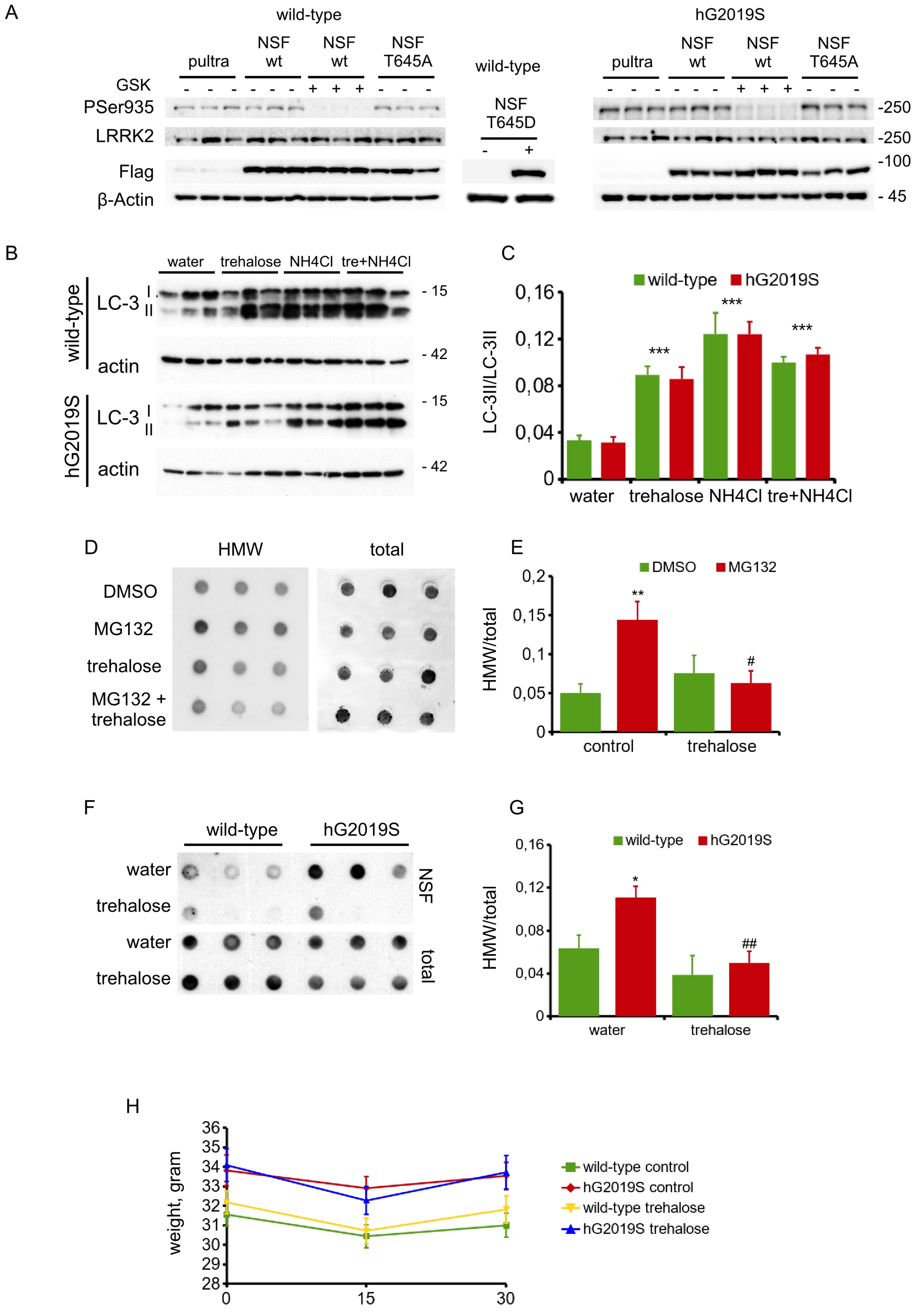


supplementary figure 6

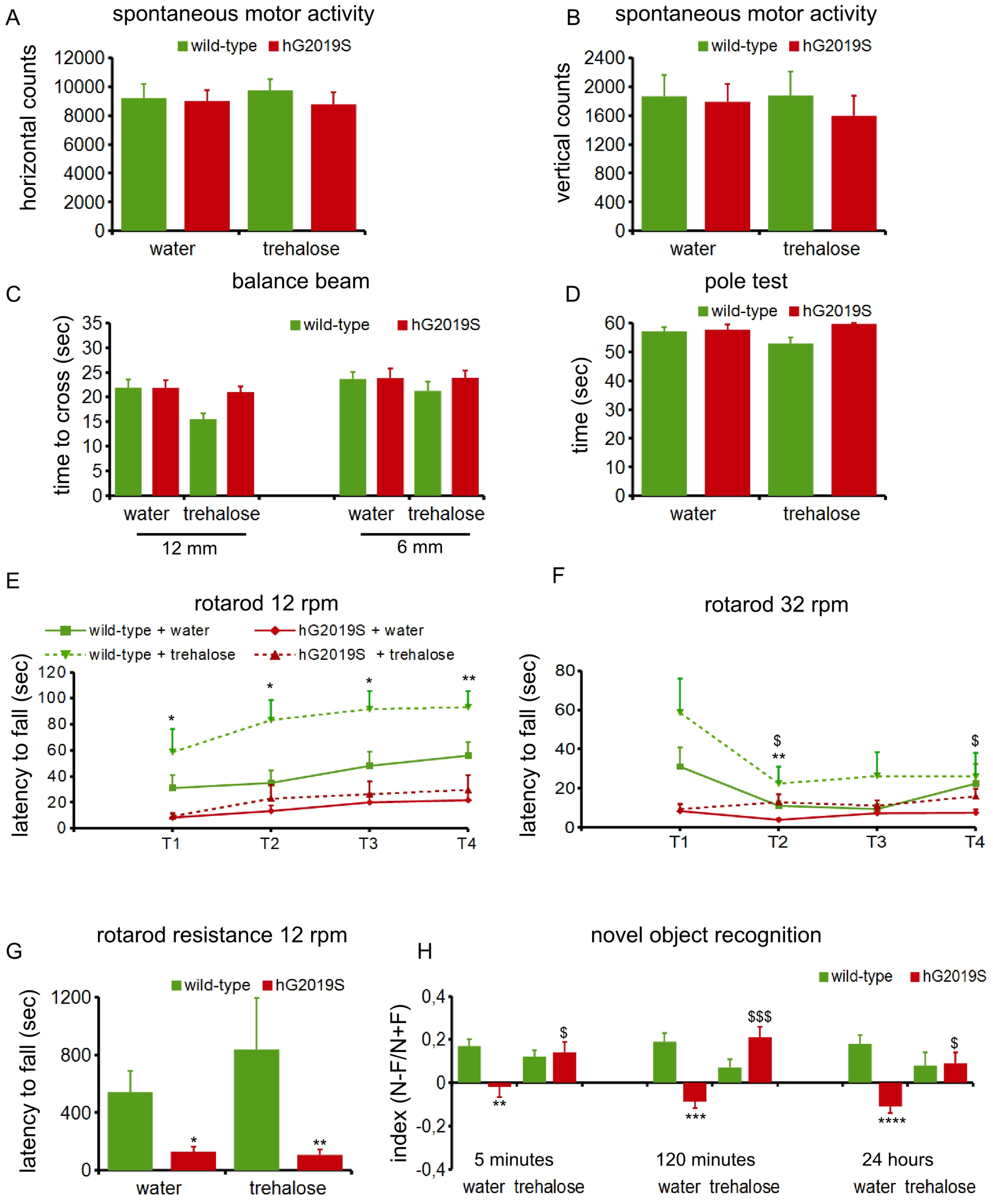


supplementary figure 7
